# Temporal single-cell atlas of full-length Huntington’s disease mouse model defines stage-specific signatures of corticostriatal dysfunction

**DOI:** 10.64898/2026.01.07.698249

**Authors:** Ashley B. Robbins, Paul T. Ranum, Icnelia Huerta-Ocampo, Michael Kuckyr, Beverly L. Davidson

## Abstract

Huntington’s disease involves progressive corticostriatal dysfunction; however, the timing of region-specific transcriptional changes remains unresolved at cellular resolution. Here, we provide a temporal single-nucleus transcriptomic atlas of striatum and motor cortex from zQ175 knock-in mice at 6 and 18 months. This full-length Huntingtin model enables staging of progressive circuit dysfunction. Striatal projection neurons show extensive early dysregulation with progressive striosomal identity erosion, whereas cortical pyramidal neuron dysfunction was layer-specific and coincided with motor symptom onset. By modeling genotype-dependent effects, we distinguish region- and cell type-specific signatures of core disease mechanisms from age-related changes and compensatory adaptations. Integrating gene co-expression and transcription factor regulatory networks, we predict candidate regulators of stage-specific dysfunction. These findings, validated across human HD and mouse model datasets, reveal temporal dynamics of disease pathogenesis in regionally distinct and interconnected neuronal populations, establishing a framework for understanding cell type-selective vulnerability.

## INTRODUCTION

Huntington’s disease (HD) is a fatal, monogenic neurodegenerative disorder caused by a CAG trinucleotide repeat expansion in exon 1 of *huntingtin* (*HTT*), producing a polyglutamine-expanded huntingtin protein^1,2^. Although mutant HTT (mHTT) is expressed ubiquitously from development through adulthood^3,4^, motor symptoms typically emerge ∼30-50 later, preceded by a prolonged prodromal phase^5–9^ characterized by asynchronous somatic CAG repeat expansion^10^. Defining the temporal sequence of molecular changes preceding cellular failure is essential for understanding selective vulnerability and identifying therapeutic targets to rescue neuronal dysfunction.

Striatal degeneration is a neuropathological hallmark of HD^11^. A 30-40% reduction in striatal volume and functional connectivity deficits can be detected a decade before clinical diagnosis^12–14^, reflecting early disruption of basal ganglia circuitry^15^. Progressive loss of spiny projection neurons (SPNs), which comprise > 90% of striatal neurons and serve as the main input of the basal ganglia motor loop, underlies striatal dysfunction^16^. Both direct (dSPN) and indirect (iSPN) pathway neurons are affected, with iSPNs exhibiting greater vulnerability^17,18^. Preferential degeneration of striosomal SPNs early in disease indicates compartment-specific susceptibility^19–21^, although the molecular basis for differential SPN susceptibility remains unclear.

HD pathology extends beyond the basal ganglia. Neuroimaging and post-mortem studies reveal disrupted corticostriatal connectivity, white matter loss, and cortical thinning^6,7,22–25^. Up to 40-50% of cortical pyramidal neurons are lost, particularly in layers III and V-VI^26,27^, and cortical atrophy correlates with motor deficit severity^28^. Recent single-cell analysis identified layer 5a as selectively vulnerable to synaptic dysfunction and degeneration in human HD^29^. Thus, early dysfunction of cortical projection neurons (CPNs) and their SPN targets implicates progressive corticostriatal disconnection in HD pathogenesis.

While cell type-specific and spatial profiling studies have advanced mechanistic understanding of HD^29–34^, temporally resolved analysis of vulnerable circuitry at cellular resolution is lacking. Addressing this gap benefits from models that allow longitudinal profiling of neurons harboring pathogenic CAG expansions over many months, allowing modeling of progressive changes.

The zQ175 knock-in model expresses full-length chimeric huntingtin with ∼188 CAG repeats under the endogenous mouse *Htt* promoter, recapitulating the genetic context and progressive neuropathology of HD^35–38^. Recent evidence suggests somatic CAG repeat expansion is associated with continuous transcriptional dysregulation prior to SPN loss^10,39^, supporting that this model effectively captures the cellular context of progressive neuronal dysfunction relevant to human disease.

Here we use a split-pool barcoding^40^ to profile cell type-specific transcriptomes from striatum and motor cortex of heterozygous zQ175 mice at 6 and 18 months, bracketing progressive cortico-striatal deficits relevant in HD^37,38,41–44^. Interrogating these regions at early and late time points enables distinction of primary pathogenic mechanisms from secondary consequences of disease. Beyond time point-specific gene dysregulation, we resolve signatures that progressively diverge from, or converge toward, physiological levels and biphasic cellular responses. We validate conservation of cell type-specific gene co-expression modules in human HD datasets and integrate our data with protein-protein interaction networks and transcriptional regulons to predict cell type- and stage-specific regulators of progressive dysfunction. Our analysis confirms and extends the characterization of sequential erosion of neuronal identity, conserved disease modules enriched for synaptic and metabolic pathways, and identifies transcriptional regulators that distinguish early adaptive responses from late pathogenic mechanisms. This resource provides a temporal framework for dissecting cell type-selective temporal vulnerability and predicts candidate drivers of cellular and circuit-level regulators of HD dysfunction.

## RESULTS

### Temporal single-nuclei profiling of the zQ175 mouse model

The zQ175 knock-in model recapitulates aspects of progressive HD neuropathology (Fig. 1a). Heterozygous mice exhibit early SPN hyperexcitability and reduced corticostriatal transmission by 3-4 months, preceding cortical pyramidal neuron (CPN) dysfunction in the motor cortex^43^.

**Figure 1.**
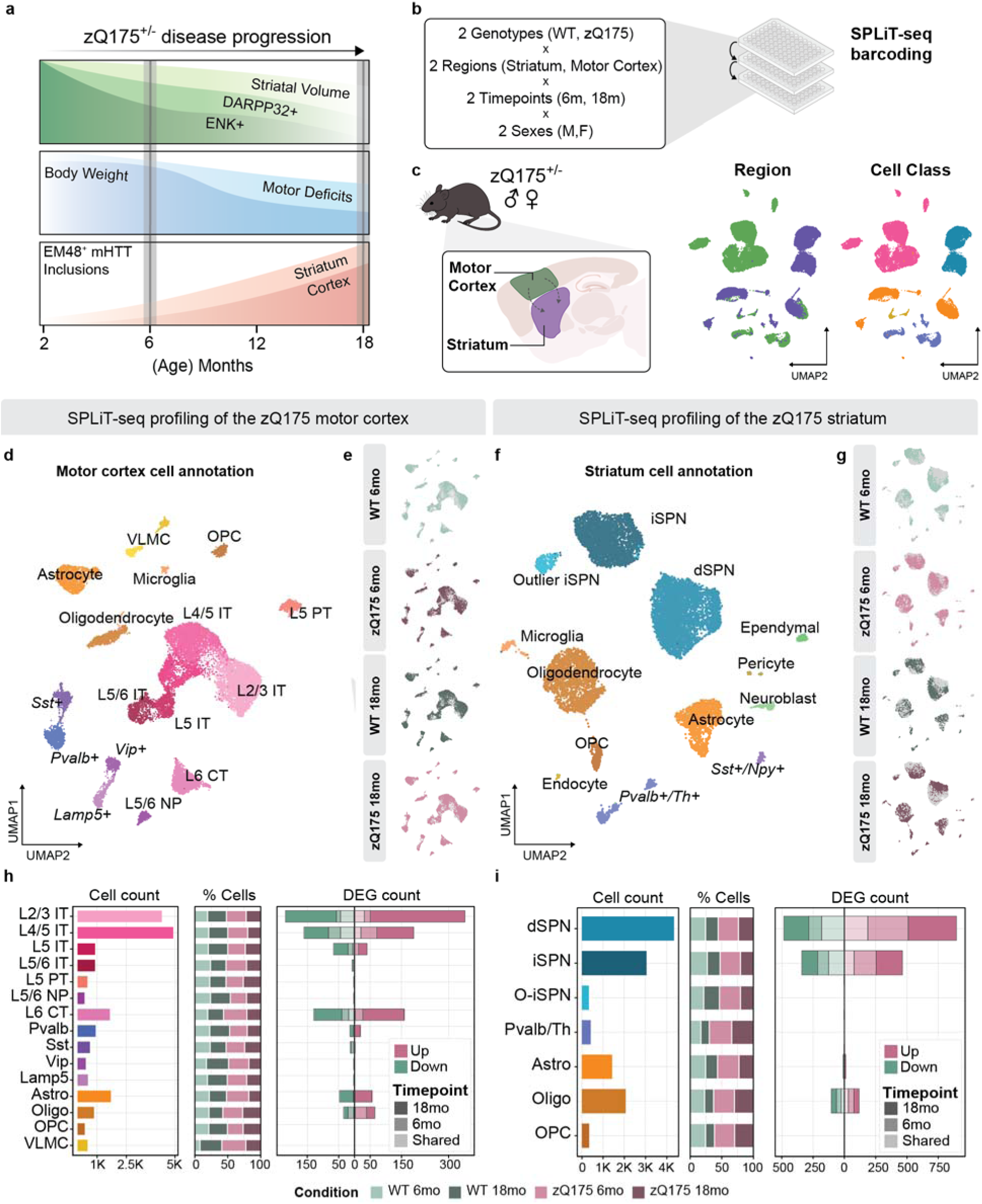
Temporal profiling of single-nuclei transcriptomes in motor cortex and striatum of zQ175 HD mice. **a** Schematic of zQ175 disease progression. **b** Experimental study design. **c** Uniform manifold approximation and projection (UMAP) plot representation of resolved single nuclei profiles colored by region and cell class. **d-g** UMAP plots of annotated cell types from motor cortex (**d-e**), and striatum (**f-g**), split by condition (**h-i**). Cell type-specific analysis of motor cortex (**h**) and striatum (**i**) displaying cell counts per cell type, cell type proportions by condition and DEGs counts separated by direction of effect (Pseudobulk limma-voom; zQ175 vs. WT; adjusted *p*-value < 0.05, |log_2_ fold change| ≥ 0.1; pink: upregulated, green: downregulated; transparency indicates temporal specificity).

Striatal volume loss is detectable by 6 months, with overt motor deficits emerging by 18 months^35–38^. We confirmed somatic CAG repeat expansion in the striatum, delayed in the cortex, and age-dependent mHTT inclusion detectable at 6 months in the striatum and 18 months in the motor cortex (Supplementary Fig. 1).

To characterize HD progression across regionally distinct cell populations, we performed single nuclei RNA-sequencing on micro-dissected striatal and motor cortex samples from zQ175 heterozygous mice and wild-type (WT) littermates at 6- and 18-month time points (Fig. 1b; WT, n = 4; zQ175, n = 4 mice per time point; 32 samples total). These time points capture cellular dysfunction before (6 months) and at the onset of motor deficits (18 months), as confirmed in our lab^37^ and others^35,45^. We optimized a combinatorial barcode-based library preparation and bioinformatics workflow to simultaneously process nuclei representing all experimental conditions in a single library^40^ (see “SPLiT-seq sublibrary generation” in Methods). Unsupervised clustering identified expected cell types in the striatum and motor cortex, with clear separation of regionally distinct cell populations and representation across experimental variables (Fig. 1c).

We separated nuclei by region to annotate cell populations (Supplementary Fig. 2a-b). CPNs were segregated into layer- and projection-specific subtypes, including intertelencephalic (IT), layer 5 pyramidal (PT), and layer 6 corticothalamic (CT) neurons (Fig. 1d and Supplementary Fig. 2a). Of note, two clusters expressing the layer 4 marker *Rorb,* consistent with Layer 4/5 IT neurons in a recent mouse primary motor cortex atlas, were combined for downstream analysis^32,46^ (Fig. 1d). Together, we identified 18 cortical and 13 striatal clusters representing major neuronal and glial cell types (Fig. 1d-g). All experimental conditions were balanced across clusters, enabling region- and cell-type specific analyses of HD-associated transcriptional changes (Supplementary Fig. 2c-f). We did not observe significant differences in cell type composition across conditions (EdgeR^47^; Supplementary Data 1).

### Regional and cell-type-specific transcriptional changes

To identify disease-associated transcriptional changes while addressing statistical challenges in single-cell data^48^, we performed differential gene expression (DGE) analysis using a pseudocell, mixed-model approach^49,50^. We compared results to pseudobulk analysis, observing highly correlated log_2_ fold change (FC) values and p-values within comparisons^51–53^. (Pearson’s r ≥ 0.9 for log_2_FC values; Spearman’s r ≥ 0.6 for most comparisons; Supplementary Data 2). The results for all time point- and cell type-specific DEG and gene set enrichment are available on Synapse (see Data Availability).

Cell type-specific DGE analysis revealed distinct vulnerability patterns across brain regions and time points. Motor cortex transcriptional dysregulation was minimal or absent at six months, with subtle changes in Layer 2-5 IT neurons, L6 CT neurons, and oligodendrocytes (Fig. 1h). However, by 18 months, cortical dysregulation was more pronounced, with L2/3 IT neurons exhibiting the most extensive changes, followed by L5 IT and L6 CT neurons. Cortical interneuron populations (Pvalb+, Sst+, Vip+, and Lamp5+) showed minimal dysregulation, though low sampling (< 5% of cortical nuclei) limited statistical power. In contrast, SPNs exhibited extensive early transcriptional changes (Fig. 1i). A high proportion of DEGs were shared across both time points, indicating sustained transcriptional perturbation, whereas 18-month DEGs were disproportionately upregulated. Oligodendrocytes in both regions displayed subtle transcriptional dysregulation at both time points (Fig. 1h-i).

These findings revealed staged neuronal dysfunction, with early striatal and delayed cortical changes. In human HD, somatic CAG expansion reaches pathogenic lengths predominantly in SPNs and CPNs, with limited expansion in glia and interneurons^10,29^. We therefore focused subsequent analyses on these cell types to investigate translationally relevant changes.

### Stage-specific transcriptional changes in striatal SPNs

To distinguish progressive disease mechanisms from static or transient responses, we modeled genotype-by-age interactions to identify genes with temporal dynamics. This approach revealed 230 genes with significant genotype-dependent temporal responses (Fig. 2a and Supplementary Data 3; |log_2_FC| ≥ 0.2; p.adj < 0.05), which we classified into three distinct temporal patterns: i) divergent from WT (Fig. 2b), ii) convergent toward WT (Fig. 2c), or iii) biphasic (Fig. 2d). We further refined this classification by applying stringent criteria to identify top genes exemplifying each pattern based on time point-specific significance and effect size (Fig. 2e; see “Pseudocell differential gene expression analysis” in Methods).

**Figure 2.**
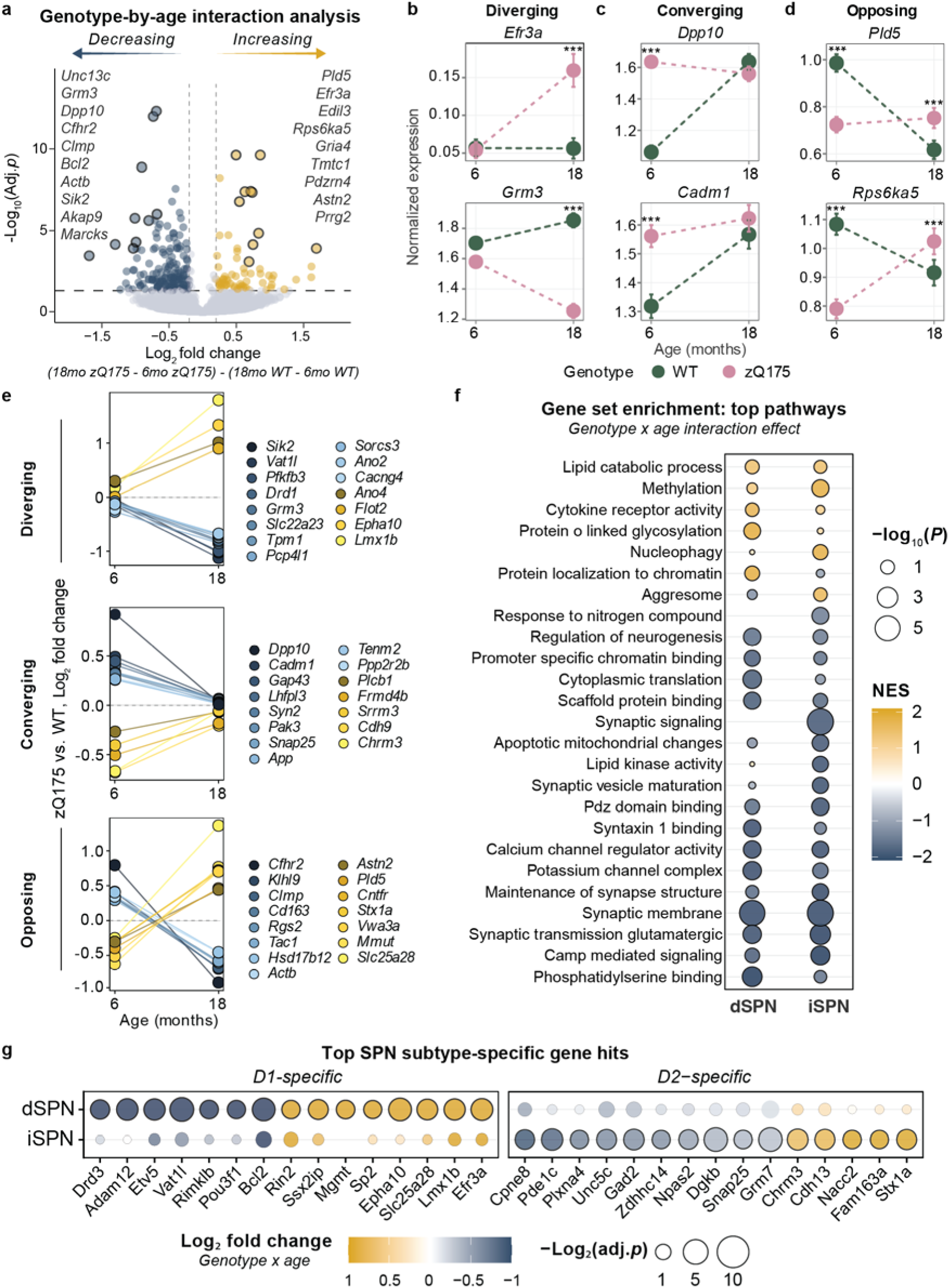
Genotype-by-age interactions define temporal expression patterns in the zQ175 striatum. **a** Volcano plot of interaction effect between zQ175 and WT from 6 to 18 months. **b-d** Average expression of representative genes with divergent (**b**), convergent (**c**), and biphasic (**d**) patterns between genotypes across time points. **e** interaction effect sizes for the top 15 genes in each pattern. **f** Top enriched pathways from GSEA analysis of genes ranked by interaction effect size. **g** Top DEGs specific to dSPN or iSPN subtypes with significant interaction effects.

*Grm3* is key example of divergent expression, encoding metabotropic glutamate receptor 3 (mGluR3). mGluR3 is involved in the regulation of presynaptic glutamate release and pharmacological rescue improves disease outcomes in HD models^54^ (Fig. 2b). Another example was *Efr3a*, involved in phosphoinositide metabolism and linked to autism spectrum disorder (ASD)^55^. Unlike *Grm3*, *Efr3a* expression increased only at 18 months (Fig. 2b). This is interesting given the negative correlation between *Efr3a* expression and BDNF-TrkB signaling observed in murine hippocampus^56^, as brain-derived neurotrophic factor (BDNF) loss is strongly linked to HD pathogenesis. Broadly, these late-diverging genes represented core processes disrupted in HD, encoding proteins regulating synaptic transmission (*Vat1l*, *Sik2*), lipid signaling (*Flot2*), glycolysis (*Pfkfb3*), and neuronal excitability (*Ano2, Ano4*, *Cacng4;* Fig. 2e).

Convergent patterns were sometimes driven by age-related WT changes, rather than zQ175 normalization. *Dpp10* and *Cadm1* exemplified this (Fig. 2c). *Dpp10,* a Foxp1 modulates voltage-gated potassium channels and neuronal excitability. We noted early elevation of *Dpp10* at 6 months concurrent with *Foxp1* downregulation (Supplementary Data 2). The human ortholog *DPP10* is downregulated in Grade 2-4 HD SPNs^30^. *Cadm1* expression encodes a cell adhesion molecule involved in synapse formation (Fig. 2c). Other top convergent genes involve synaptic vesicle release (*Snap25*, *Syn2*), synaptogenesis (*Cadm1*, *Cdh9*) and synaptic plasticity (*Gap43*, *Pak3*; Fig. 2e).

Biphasic responses were identified in synaptic plasticity (*Stx1a, Astn2, Rps6ka5* and *Rgs2*) and regulators of lipid (*Pld5*), and mitochondrial metabolism (*Mmut* and *Slc25a8*) (Fig. 2d-e). *Rps6ka5* encodes mitogen- and stress-activated protein kinase 1 (MSK-1), a BDNF-activated kinase responsible for histone H3 phosphorylation; it is elevated early then suppressed^57,58^(Fig. 2d). MSK1 levels are reduced in HD human caudate and the R6/2 mouse model, consistent with greater disease severity. Overexpression of MSK-1 confers protection against mHTT toxicity^59,60^. The expression of ciliary neurotrophic factor receptor (*Cntfr*) was also biphasic and progressively suppressed. CNTF is a trophic factor for striatal neurons with well-documented neuroprotective effects in HD^61–63^. The biphasic expression of neuroprotective signaling components suggests an early adaptive response lost over time.

Given differential vulnerability between D1-expressing dSPNs and D2-expressing iSPNs, we sought to identify subtype-specific temporal responses to chronic mHTT exposure. We performed gene set enrichment analysis, ranking genes by their age-genotype interaction effect (Fig. 2f-g, Supplementary Data 3). Both SPN subtypes showed downregulation of synaptic and glutamatergic pathways from 6 to 18 months relative to WT. However, dSPNs exhibited specific upregulation of chromatin binding and protein glycosylation pathways with dysregulation of transcription factors (*Etv5*, *Lmx1b*) and DNA repair genes (*Mgmt*). *Drd3* downregulation in dSPNs may signify loss of neuroprotective Drd3-induced mTORC1-dependent autophagy, a mechanism proposed as a therapeutic intervention to improve clearance of soluble mHTT protein^64^. In contrast, iSPNs showed signatures of metabolic stress, protein aggregation and mitochondrial dysfunction. *Nacc2* expression was upregulated in iSPNs. A transcriptional repressor involved in neurogenesis, *Nacc2* demethylation and increased expression is also predictive of Alzheimer’s diagnosis^65,66^. In agreement with dysregulation of SPN identity genes as a progressive hallmark of HD, we observed the progressive loss of dSPN marker *Pou3f1*, involved in striatal development and SPN specification^67^. We noted increased *Chrm3 expression*, a prominent iSPN marker gene that is downregulated at 6 months in both zQ175 and R6/2 mice^30^. These subtype-specific temporal signatures confirm known differential vulnerability of iSPNs versus dSPNs while revealing distinct transcriptional trajectories. dSPNs exhibit progressive transcriptional and epigenetic dysregulation, whereas iSPNs show earlier metabolic stress signatures. Notably, biphasic expression patterns across both subtypes distinguish genes with transient versus sustained dysregulation as disease progresses.

### Temporal progression of striatal SPN identity loss in HD

In addition to direct-indirect pathway division, SPNs are organized in distinct striatal compartments (Striosome, Matrix; Fig. 3a). Recent studies demonstrated preferential depletion of striosomal SPNs in Grade 0-1 HD preceded by an erosion of transcriptional distinction between compartmental identities^20,21^. Furthermore, sustained loss of dopamine receptors and key signaling molecules, including phosphodiesterase 10A (PDE10A) a critical cAMP regulator in SPNs, has been noted in striosomal zQ175 SPNs by 3 months of age^19^. However, temporal progression of identity loss remains unexplored.

**Figure 3.**
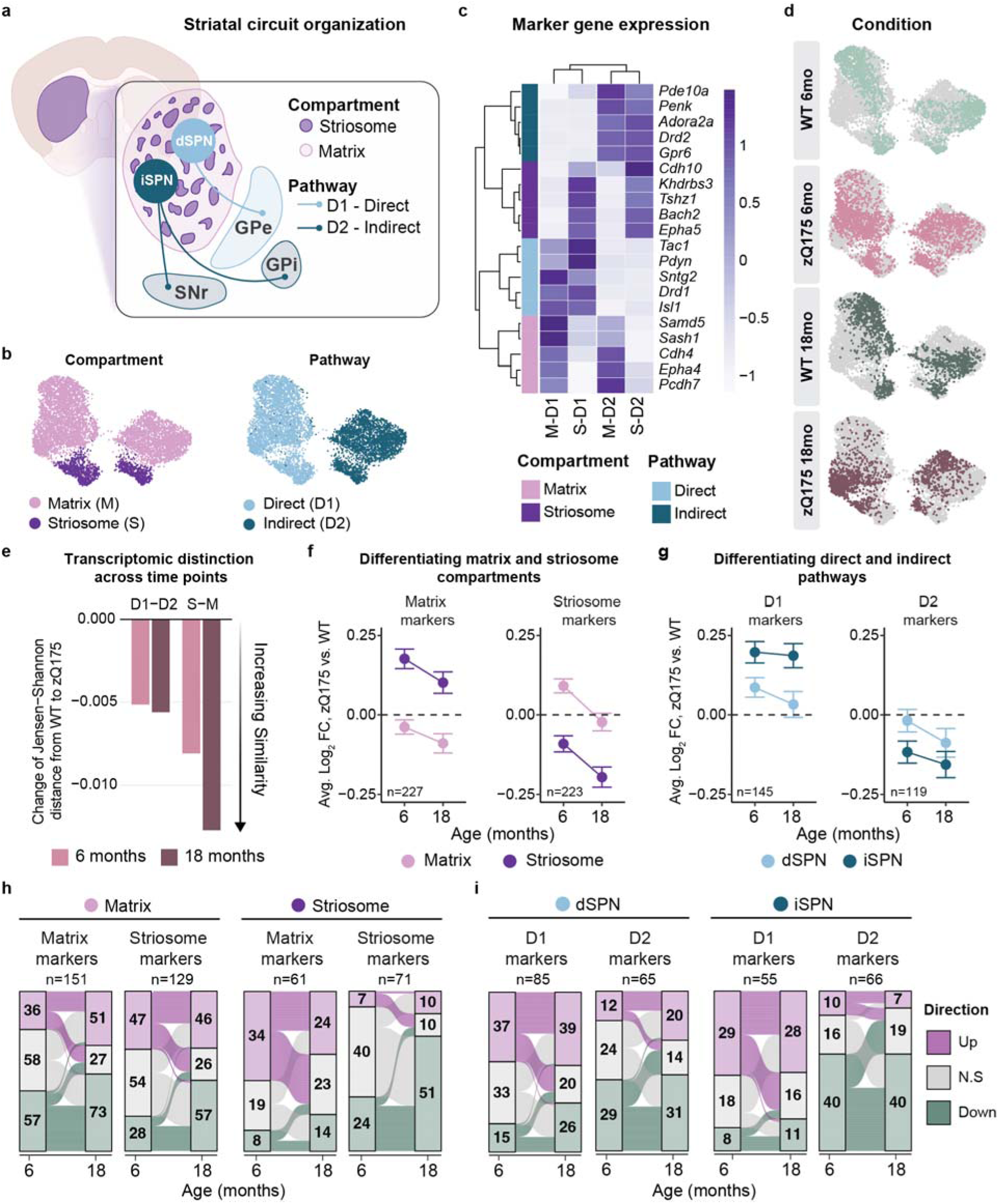
Progressive loss of compartment and pathway identity in zQ175 SPNs. **a** Schematic of striatal organization. Gpe, external globus pallidus; GPi, internal globus pallidus; SNr, substantia nigra pars reticulata. **b** UMAP plots colored by SPN identity. **c** Scaled expression of canonical pathway and compartment gene markers across SPN subclusters. **d** UMAP plot of the SPN dataset split by condition. **e** Jensen-Shannon distance analysis showing the loss of transcriptomic distinctions between striatal subtypes. Comparisons are shown separately between dSPN (D1) and iSPN (D2), then striosome (S) and matrix (M). **f** Average log_2_ fold change values of matrix and striosomal markers by compartment identity at 6 and 18 months and **g** D1 and D2 markers by pathway identity at 6 and 18 months. Error bars showing ± standard error of the mean (s.e.m). The total number of genes in each signature are noted in each panel; n = 227 Matrix marker genes; n = 223 Striosome marker genes; n = 145 D1 marker genes; n = 119 D2 marker genes. **h-i** River plots quantifying the number of dysregulated D1 and D2 markers at 6 and 18 months for each pathway (dSPN, iSPN).

To address this, we subclustered and annotated SPNs by compartment (Matrix, Striosome) and pathway (D1-direct, D2-indirect) identities, with balanced representation across genotypes and time points (Fig. 3b-d). We quantified transcriptional distinction between these populations using Jensen-Shannon distance and observed an increased similarity (identity erosion) between both compartment (S-M) and pathway (D1-D2) SPNs in zQ175 compared to WT SPNs at each time point (Fig. 3e). Our 6-month data is consistent with earlier reports in 6-month zQ175 and R6/2 transgenic mice^19^. Additionally, we identify progressive dysregulation of identity genes distinguishing S-M compartments at 18 months. In contrast, D1-D2 distinction remained relatively stable across time.

To examine compartmental identity in detail, we define striosome and matrix enriched genes (Supplementary Data 4; |log_2_FC| ≥ 0.2 and p.adj < 0.001). At 6 months, compartment-specific marker genes were upregulated in the opposing compartment, such that striosomal SPNs upregulated matrix identity genes and matrix SPNs upregulated striosomal identity genes (Fig. 3f). Downregulation of matrix and striosomal identity genes in their respective subpopulations occurred at both time points. By 18 months, marker expression decreased in both compartments, most pronounced for striosomal markers in striosomal SPNs (Fig. 3f). Pathway marker gene dysregulation was less dynamic (Fig. 3g), though dSPNs trended toward greater loss of identity gene expression over time.

Quantification confirmed these patterns (Fig. 3h-i and Supplementary Data 4; |log_2_FC| ≥ 0.1; p.adj < 0.05). The total number of dysregulated matrix and striosomal identity genes increased from 6 to 18 months in matrix SPNs (Fig. 3h). Contrastingly, striosomal SPNs exhibited a loss of previously upregulated matrix identity genes, as reflected by the loss of distinction from WT expression (Fig. 3f). This was concurrent with doubling of downregulated striosomal identity genes (Fig. 3h, right panel; n = 24, 6 months; n = 51, 18 months). Pathway-specific identity gene dysregulation suggested greater stability of the D1-D2 axis over time, while the underlying signatures again demonstrated both stable and dynamic changes (Fig. 3i).

We narrowed our analysis to compartment, and pathway identity genes conserved in human SPNs (Supplementary Fig. 3a, c)^20^. Over half in each set (56.7 - 63.7%) were also significantly dysregulated in zQ175 SPNs. Temporal analysis (Δ log_2_FC = (18 month, zQ175 vs. WT) - (6 month, zQ175 vs. WT)) highlighted both time point-specific and progressive signatures (Supplementary Fig. 3b, d). Matrix SPNs exhibited upregulated expression of several conserved striosomal markers at 6 months (e.g., *Oprm1*, *Magi2*, *Sema6d*), with few changes specific to 18 months. Striosomal marker *Lin28b*, a developmental regulator of neurogenesis and cellular differentiation^68,69^, was among the genes upregulated at 18 months in matrix SPNs, supporting reactivation of developmental programs following progressive loss of striosome-matrix boundary maintenance (*Epha4*, *Epha5*)(Supplementary Fig. 3a-b). Among conserved D1 and D2 pathway markers, D2 markers *Penk*, *Drd2* and *Adora2a* were downregulated at 6 months, consistent with previous findings^20,30,70^, while D1 marker dysregulation was largely delayed (e.g.,*Tac1*, *Drd1*)(Fig. 3h and Supplementary Fig. 3c-d). Transient changes at 6 months included those involved in lipid metabolism (*Osbpl10*), cellular stress (*Ola1*), and signal transduction (*Prkca*, *Gpr139*, *Pde4d*). Collectively, our data confirm compartment-specific vulnerability in HD^20,21^ and reveal progressive erosion of both striosome-matrix compartmental boundaries and pathway-specific molecular identities with distinct temporal trajectories.

### Genotype-associated modules in SPN co-expression networks

To systematically identify coordinated transcriptional changes in SPNs, we performed weighted gene co-expression network analysis (hdWGCNA)^71^. This approach enabled identification of disease modules with temporal dynamics and cell type-specificity, further resolving gene networks identified with bulk tissue analyses^72^. We identified 12 gene co-expression modules, encompassing distinct expression patterns across SPN subtypes, genotypes, and time points (Fig. 4a-b and Supplementary Data 5). These modules captured distinct biological processes, including RNA splicing and glucose homeostasis (SPN-M2), synaptic signaling and maintenance (SPN-M4, M6, and M10), proteostasis (SPN-M5), metabolic stress (M7), and neurodevelopmental programs (SPN-M3, M11, and M12) (Supplementary Fig. 4a).

**Figure 4.**
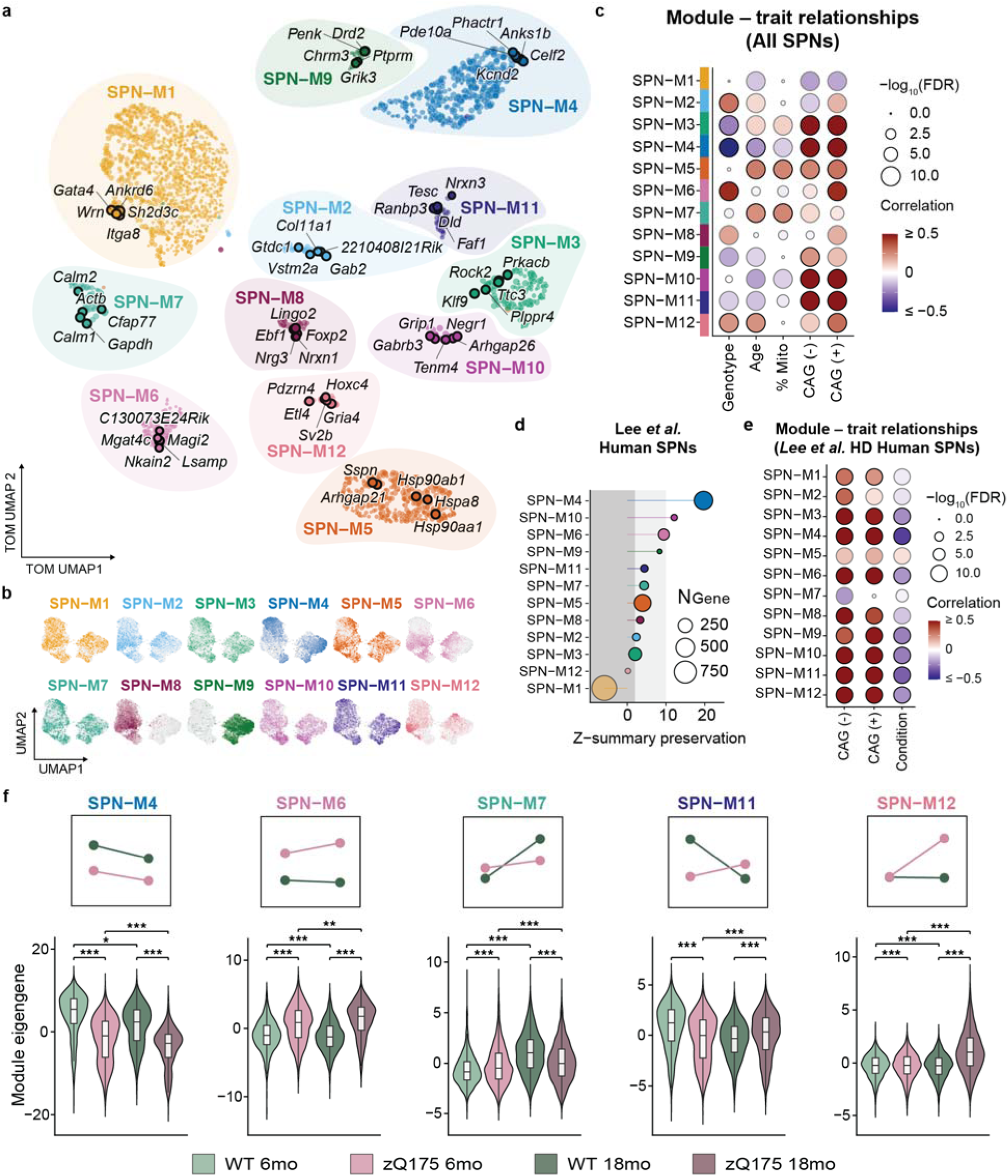
SPN gene modules reveal stage-specific dysregulation and opposing molecular aging trajectories in HD. **a** Supervised UMAP plot of the SPN co-expression network. Each node represents a gene, colored by module assignment and scaled by kME. The top 5 hub genes ranked by kME are annotated for each module. **b** UMAP plots of SPN nuclei colored by each module eigengene (ME). **c** Module-trait relationships represented by the Pearson correlation for each module eigengene and covariate. CAG (+) and CAG (-) scores with the UCell algorithm. % Mito: Mitochondrial RNA reads. Correlation coefficients for genotype reflect expression in zQ175 vs. WT SPNs. **d-e** Analysis of SPN hdWGCNA module preservation in the human HD SPNs, data from Lee et al.^30^ (**d**) and R6/2 mouse model, Lim et al.^73^ (**e**) *Z* > 10, highly preserved; *Z* > 2, moderately preserved, and *Z* < 2, not preserved. Correlation coefficients for condition in (**e**) reflect expression in HD vs. Control. **f** Module expression scores by experimental condition. A linear model was fit for each subtype and time point, accounting for covariates of library batch and sex, to determine genotype and age effect at each time point (6 and 18 months). Insets show mean expression at each time point by genotype. \**p* < 0.05, \*\**p* < 0.001, \*\*\**p* < 0.0001.

We prioritized modules by correlation with i) genotype, ii) age, and iii) CAG repeat length-dependent gene signatures^10^ (Fig. 4c). Five modules were associated with genotype (SPN-M2, M3, M4, M6, and M12; |*r|* > 0.2, adj. *p* < 0.0001) and four with age (SPN-M4, M5, M7, and M12). SPN-M3-6 and SPN-M12 were also positively correlated with the expression of CAG repeat-dependent genes (Fig. 4c and Supplementary Fig. 4b). SPN-M5 and SPN-M7 both correlated with age and were enriched for genes in the CAG repeat-independent “discrete” HD gene signature identified in human SPNs^10^ (Fig. 4b; 719 genes).

Most modules demonstrated cross-species and cross-model preservation (Z > 2) with a human HD SPN datasets^30^ and the R6/2 mouse model data^73^ (Fig. 4d and Supplementary 4d). SPN-M11 was moderately preserved in human SPNs (*Z* = 4.43), but not in R6/2 (Z < 2, R6/2). SPN-M4, M6, M7 and M11 represent distinct temporal dynamics via sustained (e.g., SPN-M4, M6) or biphasic (e.g., SPN-M7, SPN-M11) expression across time points (Fig. 4f and Supplementary Fig. 4f-g).

Module SPN-M4 exhibited the strongest disease association (genotype, *r* = -0.51; age, *r* = -0.21; Fig. 4c), with progressive downregulation at 6 and 18 months (Fig. 4f, Supplementary Fig. 4d). This module involved glutamatergic synaptic transmission, calcium regulation, and GTPase signaling pathways (Supplementary Fig. 4a), converging onto the mGluR1-PLC-IP3R1 signaling cascade, including receptors (*Grm1, Gpr158*), signal transducers (*Plcb1, Itpr1, Dgkb, Dgki*), effectors (*Prkcb, Cacna2d3, Pde10a*), and downstream modulators (*Rgs9, Rbfox1, Celf2*) (Fig. 4a and Supplementary Fig. 6). Following integration of the gene co-expression network with functional protein interaction networks from a public database (STRING^74^), we observed a large overlap of hub genes based on gene module (kME) and protein-protein interaction connectivity further supporting the convergence of M4 on proteins interacting this signaling cascade (Supplementary Fig.5a). Furthermore, there was a significant enrichment of HTT interacting proteins encoded in the top 25 hub genes (e.g., *Itpr1, Pde10a, Dgkb, Gsk3b*; Supplementary Fig. 5b-c and Supplementary Data 5).

M4 was highly preserved in HD patient and R6/2 SPNs, correlated negatively with disease (Fig. 4d-e and Supplementary Fig. 4d-e) and showing consistent low expression relative to WT (Supplementary Fig. 4f). This module contains many genes previously associated with CAG repeat length in both human post-mortem datasets and allelic mouse series^10,72^, converging most specifically onto the conserved mGluR1-PLC-IP3R1 signaling pathway.

To identify the potential drivers of progressive dysregulation in M4 we identified top hub genes with time point-restricted or significant interaction effects (Supplementary Fig. 6). *Zswim6*, a repressive epigenetic regulator associated with SUZ12, a core component of the Polycomb Repressive Complex 2 (PRC2), emerged as a hub gene with a significant interaction between genotype and age (log_2_FC = -0.27, adj. p = 0.02). *Zswim6* has recently been implicated in corticostriatal synapse development, specifically via the regulation of chromatin accessibility and transcriptional dysregulation^75^. Additional top hub genes with progressive changes included RNA-binding protein-encoding genes *Rbfox1*^76,77^ and *Celf2*^78^, which both regulate calcium channel and synaptic protein expression through alternative splicing. M4 confirms mGluR1-PLC-IP3R1 signaling pathway dysregulation as a central mechanism of SPN dysfunction in HD, with our temporal analysis revealing progressive collapse across SPN subtypes and predicting roles of network hub genes involved both epigenetic (Zswim6) and post-transcriptional (Rbfox1, Celf2) regulation.

On the other hand, SPN-M6 showed sustained upregulation in zQ175 SPNs (Fig. 4f and Supplementary Fig. 4f), top hub genes involved in synaptic transmission and ion homeostasis (e.g., Celf4, Gria4, Gabrb1, Nkain2, Cadm1) exhibited early elevation that normalized by 18 months (Supplementary Fig. 6), suggesting stage-specific transient responses. This contrasted with modules showing sustained responses (SPN-M2: splicing/proteostasis; SPN-M3: activity-dependent genes), where dysregulation persisted across timepoints (Supplementary Fig. 4f and Supplementary Fig.6).

**Figure 5.**
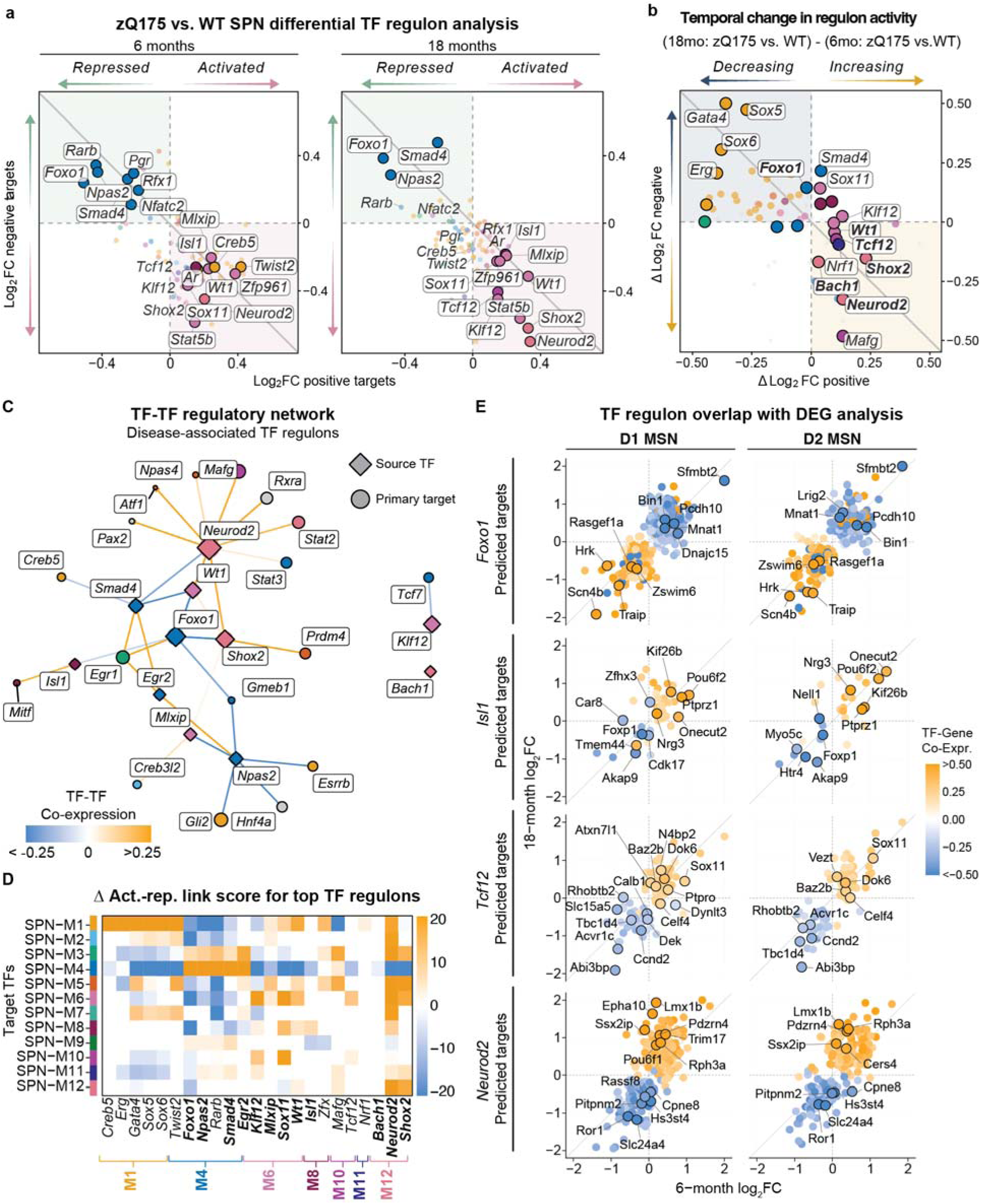
Transcription factor regulatory network dysregulation in zQ175 SPNs. **a** Differential TF regulon analysis in zQ175 vs. WT SPNs at 6 months and 18 months. Scatter plots show log_2_ fold change values for negatively correlated genes (x-axis) vs. positively correlated genes (y-axis) of significant TF regulons. Upper left quadrant: repressed TF regulons; Bottom right quadrant: activated TF regulons. **b** Temporal progression of regulon dysregulation, represented by the change in effect size from 6 to 18 months in zQ175 SPNs relative to WT. Upper left quadrant: TF regulon activity decreased over time; Bottom right quadrant: TF regulon activity increased over time. **c** TF-TF regulatory network plot depicting regulatory links between the top dysregulation regulon TFs, shared across both time points and their primary downstream TF targets. Nodes represent individual TFs colored by module identity and size depicting the degree of co-expression linkages within the network. Edges reflect the Pearson correlation coefficient of co-expression between TFs above a threshold of 0.1. **d** Heatmap showing the difference in interaction strength between activating and repressing TF-gene correlations of individual TFs and their downstream primary targets within SPN co-expression modules. Top TFs shared across both time points are bolded. **e** Overlap of predicted target genes in select top dysregulated regulons and time point-specific zQ175 vs. WT DEGs split by SPN pathway identity. Color depicts the correlation coefficient between each TF and individual genes. Labeled genes reflected DEGs with the greatest regulatory score or the importance of that TF for modeling gene expression of a given gene.

**Figure 6.**
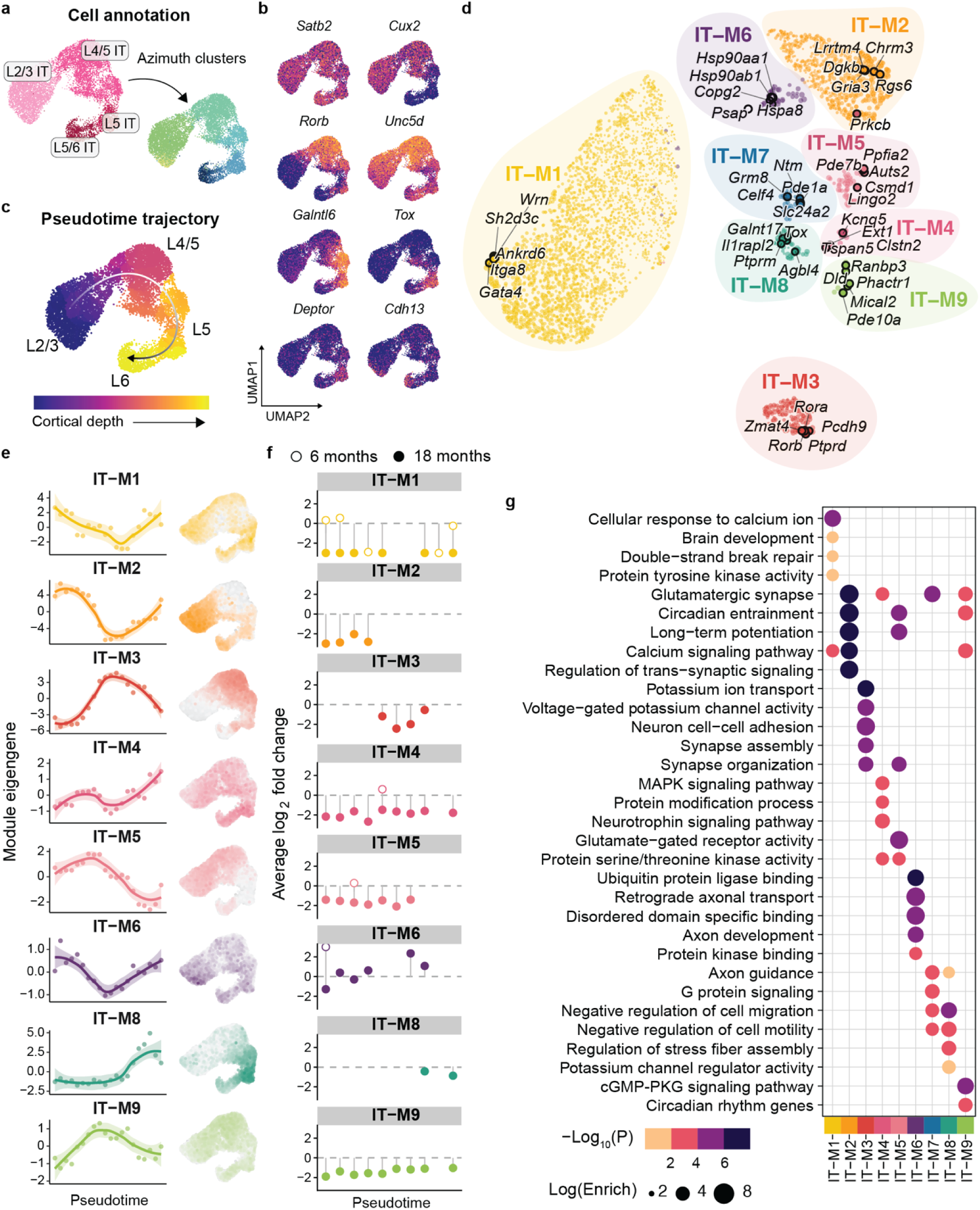
Layer-specific hdWGCNA modules capture disease signatures in zQ175 IT neurons. **a** UMAP of layer-specific annotations in IT neuron subset by unsupervised clustering and reference mapping. **b** Representative layer-specific marker genes for cortical layers in IT neurons. **c** UMAP of pseudotime trajectory in IT neurons. **d** Supervised UMAP plot of the IT co-expression network. Each node represents a gene, colored by module assignment and scaled by kME. Top 5 hub genes are annotated. **e** Module eigengene expression plotted as a function of pseudo trajectory. Cells were ordered by cortical layer from L2/3 to L6 via unsupervised clustering, in agreement with reference-based annotation. UMAP plots depicting single nuclei eigengene values are also displayed. **f** Differential module eigengene analysis across cortical layers L2/3 to L6 at 6- and 18-month time points. Nuclei were segregated into 25 bins based on pseudo trajectory, reflecting cortical layer depth. **g** Dot plot of select GO and KEGG pathway enrichment distinguishing co-expression modules.

Two modules revealed biphasic temporal patterns that distinguish transient cellular responses from sustained disease signatures (SPN-M7 and SPN-M11; Supplementary Fig. 4g). SPN-M7 exhibited biphasic regulation, elevated at six months, and downregulated at 18 months in Q175 SPNs. This was in contrast to WT, where M7 showed a progressive age-dependent increase (Fig. 4f and Supplementary Fig. 4g). Investigation of temporally dynamic hub genes revealed that M7 is a cellular stress response signature, including primary calcium sensors (*Calm1*, *Calm2*), proteostasis machinery (*Ubb*, *Ppia*, *Psap*), mitochondrial regulators (*Ghitm*, *Amd1*, *Amd2*), and stress-responsive glycolytic enzyme encoded by *Gadph*^79^ (Supplementary Fig. 6). The levels of nucleolar protein nucleolin, encoded by the hub gene *Ncl*, correlated with the levels of nuclear mHTT inclusions in R6/2 mice nuclei and a predicted HTT interactor (Supplementary Fig. 5a), further implicating this module in stress-related responses to mHtt aggregation^80^

Our data suggest that the early upregulation of M7 reflects an adaptive stress response that is no longer dynamic to age-related changes in zQ175 SPNs by 18 months. In the R6/2 model, in which nuclear aggregation of mHTT is an early event, there is a positive correlation with genotype, whereas in the human SPN dataset, there is no correlation with disease status. The M7 module had the greatest overlap to the “discrete” gene signature in Handsaker et al.^10^, which is upregulated in SPNs with long CAG repeats (> 150). Together, these findings define M7 as a stress response signature that is transiently elevated early but not sustained, distinguishing this potentially adaptive response in the zQ175 knock-in model from sustained dysregulation in a transgenic model (R6/2).

SPN-M11 showed complex age-dependent changes in the expression of synaptic maintenance genes (*Pclo*, *Nrxn3, Tesc*; Supplementary Fig. 4e and Supplementary Fig. 6). Unlike the biphasic stress response captured by SPN-M7, M11 reflects an age-dependent decline in genes involved in synaptic maintenance and remodeling. This pattern is preserved in human HD SPNS, but not in R6/2, where M11 expression is positively correlated with genotype but negatively with CAG repeat dependent gene expression (Fig. 4d-e and Supplementary Fig. 4d-e).

Unlike other disease-associated modules, SPN-M12 exhibited a late-emergent pattern, expressed predominantly at 18 months in zQ175 SPNs with minimal expression at 6 months or in WT SPNs (Fig. 4f). Hub genes included developmental transcription factors normally silenced in mature SPNs (e.g., *Hoxc4, Hoxc5*, *Pou6f1*, *Pknox2*, and *Shox2*). Of note, *Hoxc4* and *Hoxc5* upregulation is seen in human SPNs as part of a developmental “de-repression” gene signature^10^ correlated with highly expanded CAG repeats. SPN-M12 positively correlated with CAG repeat signatures and was detected in less than 0.5% and 5% of WT and zQ175 SPNs, respectively.

Beyond developmental factors, M12 hub genes represented specific disruption of components involved in synaptic vesicle release. Synaptic vesicle glycoprotein 2 (*Sv2b*), crucial for neurotransmitter release, was recently identified as a potential genetic modifier of HD onset^81^ and whose expression increased with CAG repeat length in human SPNs^10^. Additional hub genes were also involved in the synaptic vesicle cycle, including *Kif1b* responsible for the trafficking of synaptic vesicle precursors and mitochondria to axon terminals and *Rph3a* mediating vesicle docking and fusion. Collectively, SPN-M12 represents a late-emerging transcriptional state marked by transcriptional de-repression and dysregulation of synaptic vesicle dynamics critical for neurotransmitter release.

Collectively, these twelve modules confirm core HD pathways, while importantly revealing temporal dynamics that distinguish: (i) sustained pathogenic dysregulation (M4, M6), (ii) transient biphasic responses (M7, M11), and (iii) late-emergent de-repression (M12). Cross-species preservation and corroboration in human CAG repeat-length dependent signatures validates the relevance of these modules to human disease, while hub gene analysis informed candidate regulators for future mechanistic studies.

### Stage-specific TF regulatory network analysis in HD SPNs

To identity transcriptional regulators driving the temporal patterns observed in our module analysis, we defined transcription factor regulatory networks, linking transcription factors to downstream target gene co-expression (Supplementary Figure 7a; see “Transcription regulatory network and regulon analysis” in Methods)^71,82^.

We identified 34 TF regulons with differential activities between zQ175 and WT SPNs (Supplementary Data 6; |average log_2_FC| > 0.1 & adj. *p* < 0.05) (Fig. 5a-b). Twelve TF regulons showed persistent dysregulation across both time points, suggesting core transcriptional disturbances. These included TFs associated with HD relevant modules: SPN-M4 (*Foxo1*, *Npas2*, *Smad4, Egr2*), SPN-M6 (*Klf12, Mlxip, Wt1*), and SPN-M12 (*Bach1, Shox2, Neurod2*) (Fig. 5c-d), all genotype- and age-dependent modules enriched in CAG repeat length-dependent genes.

Foxo1, a forkhead box transcription factor activating oxidate stress responses and autophagy, showed decreased activity in zQ175 SPNs, consistent with impaired cellular stress adaption in HD (Fig. 5e and Supplementary Fig. 7b-c). Other repressed TF regulons in SPN-M4 included Npas2 (a circadian clock component), Smad4 (a TGF- β/BMP signaling mediator), and transcription factor Egr2. Together, we observed repression of regulatory activity for TFs in SPN-M4, consistent with transcriptional depression of core SPN processes in HD (Fig. 5d). Analysis of predicted Foxo1 target genes revealed progressive dysregulation across both dSPN and iSPNs, with 33% of predicted regulon targets showing significant differential expression (Fig. 5e) (e.g., *Zswim6, Scn4b, Traip, Sfmbt2*).

Isl1 regulatory activity was increased in zQ175 SPNs across timepoints and SPN subclasses, with the exception of 18-month Drd1^+^ striosomal SPNs (Fig. 5a-b and Supplementary Fig. 7b). Associated with the dSPN identity module SPN-M8, Isl1 is required for the specification and maturation of SPNs^83,84^. Interestingly, Isl1 has also been identified as an upstream transcriptional regulator of *Foxo1* during striatal development. In *Isl1*^-/-^ murine SPNs, the loss of *Foxo1* expression was linked to reduced DARPP-32 levels and dSPN survival^83^. In the context of mHTT, Foxo1 confers neuroprotection by reducing the accumulation of toxic protein aggregates and aberrant RNA species by decreasing both the level of expanded CAG transcript- and protein levels^85,86^. Our analysis supports Foxo1 as a possible direct regulator of *Isl1* expression that is impacted in HD; there are Foxo1 responsive elements in the *Isl1* locus^87^,and loss of Isl1 regulation of the canonical dSPN identity gene *Nrg3* (Neuroregulin 3), other transcription factors (e.g., *Onecut2, Pou6f2),* chromatin regulators (e.g., *Phf7, Zhx3*), and the sumoylation modifier *Pias1*^88^ (Fig. 5e and Supplementary Fig. 7d). Additionally, *Foxp1* expression is predicted to be repressed by Isl1 activity, pointing to dysregulation of the Foxo1-Isl1 regulatory network in the progressive remodeling of dSPNs toward a more immature cell state (Fig. 5c).

Another predicted TF target of Foxo1 was *Wt1* encoding Wilms’ tumour 1, associated with SPN-M6 and identified as a predicted regulator of activated gene signatures in human HD SPNs^30^ (Fig. 5a-d). Wt1 regulates DNA methytransferase 3A (DNMT3A), influencing genome-wide DNA methylation at promoters and implicated in HD epigenetic mechanisms^89,90^. Increased Wt1 activity, as seen in zQ175 SPNs, results in widespread transcriptional repression^87^. Analysis of the Wt1 regulatory network in our model reveals a number of SPN-M4 hub genes as top targets of Wt1 activity induced repression (e.g., *Pde10a, Pde1b, Kcnd2, Adcy5, Anks1b, Phactr1, Zswim6*), whereas TF *Neurod2* expression was positively correlated (Supplementary Fig. 7b, e). Other TFs associated with SPN-M6 with increased activity include BDNF regulator Klf12^91^ and glucose metabolism regulator Mlxip^92^. Module-level regulatory analysis revealed SPN-M4 module genes were enriched as negative downstream targets of SPN-M6 TFs.

TFs in SPN-M12 also exhibited positive differential activity in zQ175 SPNs. *Shox2, Neurod2* and *Bach1* are the most progressive activated regulons in our analysis. Neurod2 was upstream of several disease-relevant TFs (Fig. 5c) including those involved in neurodevelopment (*Shox2* and *Pax2*) and DNA methylation (*Wt1*, *Atf3*), Moreover, Neurod2 was a predicted repressor of striatal neuronal survival mediator *Foxo1*, potentially driving progressive loss of Foxo1 activity in Q175 SPNs (Fig. 5c). Importantly, *Neurod2* expression is increased in HD SPNs^10^. Our data suggests that Neurod2 activity is altered with disease progression and may contribute to neurodevelopmental reprogramming and SPN dysfunction in the striatum.

SPN-M10 *Tcf12* and *Sox11* TF regulons were activated at 18 months and unchanged earlier. Tcf12 was previously identified as a regulatory transcription factor^93^ affecting histone acetylation in HD by a meta-analysis of novel HD gene variants. More broadly, Tcf12 functions in neuronal development via chromatin remodeling^94^. Sox11 is also involved in neurodevelopment and neuronal survival and a predicted target of Tcf12 (Fig. 5e).

In sum, by integrating differential gene expression analysis, co-expression modules and TF regulatory networks, we identify multiple interconnected transcriptional programs, demonstrate temporal and module-specific activity patterns in the zQ175 model, and define predicted regulatory associations between human HD-relevant TFs.

### Layer-dependent transcriptional changes in IT neurons

We observed few early changes but delayed transcriptional signatures at 18 months in CPNs (Fig. 1h), unlike striatum. To capture progressive gene signatures involved in corticostriatal dysfunction, we performed hdWGCNA analysis on intertelencephalic (IT) pyramidal neurons, which send their projections to contralateral direct pathway SPNs and cortical targets^46^. Nine gene co-expression modules were defined in IT neurons (Supplementary Fig. 8).

IT neurons form a continuous spectrum of gene expression that correlates with cortical depth^32^. To assess module eigengene expression across cortical layers, we first defined subpopulations by reference-based annotation (Fig. 6a), then utilized pseudotime trajectory analysis to order cells cortical depth (Fig. 6b), consistent with the expression of canonical marker genes (Fig. 6c). Assessment of module eigengene expression revealed progressive loss of gene expression by 18 months, associated with cortical layer identity and distinct biological processes (Fig. 6e-g).

IT-M2, M3, M5 and M8 were enriched in layer-specific subpopulations (Fig. 6f). We assessed differential expression of top hub genes in these modules in discrete subclusters and identified progressive layer-specific genes dysregulation of subclass-specific markers (Supplementary Fig. 10a). Layer 2/3 marker gene *Sorcs3* was downregulated by 18 months, whereas *Gpc6* was upregulated in both L2/3 and L5 IT neurons. Layer 5 marker genes *Hs6st3* and *Cdh12* (IT-M3) were also downregulated by 18 months, alongside Layer 5a marker *Deptor* (IT-M8) (Supplementary Fig. 8a).

Additional modules were expressed across all IT neurons (IT-M1, IT-M4, IT-M6, IT-M9; Fig. 6e). IT-M7 was the only module with no disease-associated expression likely due to limited sampling of layer 5b-6 CPNs, in which this module was enriched.

Module IT-M4 was globally dysregulated, enriched for signal transduction, serine/threonine kinase activity and synaptic function (Fig. 6g). Protein-protein interaction network analysis refined our hub genes to a central synaptic transmission network (STRING^74^, Fig. 7a). *Homer1*, a central hub and predicted HTT interactor^74^, is highly connected to other synaptic genes (*Nrcam*, *Prkcb*, and *Snap25*) and critical for activity-dependent plasticity and BDNF signaling. *Ntrk2*, encoding the BDNF receptor TrkB, was another top hit predicted to interact with Homer1. The coordinated downregulation of *Ntrk2* and *Homer1* at 18 months suggests disrupted BDNF signaling by 18 months (Fig. 7b). Overlap with human HD gene signatures in motor cortex CPNs^29^ revealed conserved downregulation of module genes in IT-M4 (Fig. 7c and Supplementary 8c), including *HOMER1*, *GABRB1*, and *SNAP25* (Supplementary Data 7).

**Figure 7.**
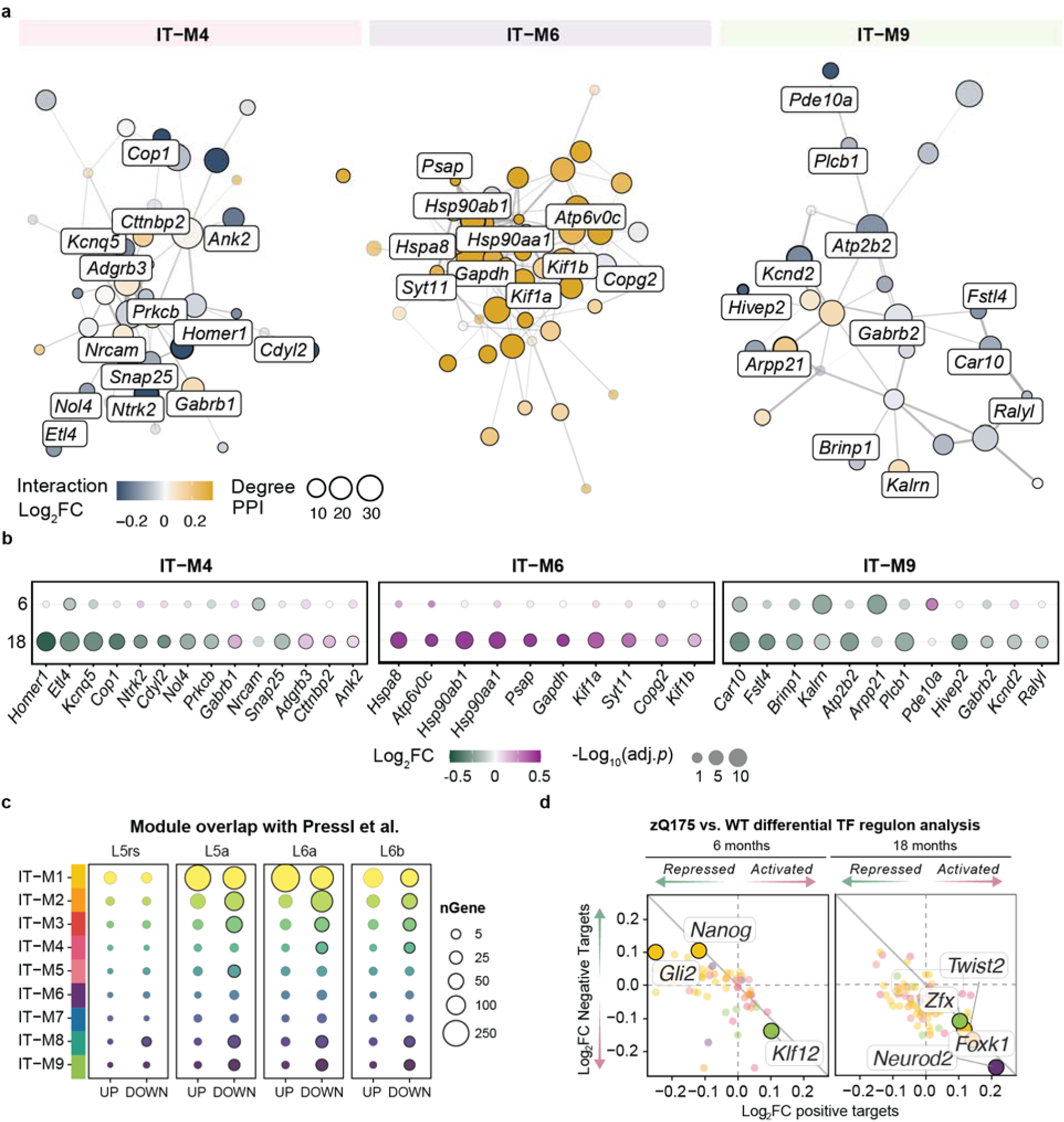
Late emergent disease signatures in motor cortex IT neurons. **a** Integrated gene co-expression and protein-protein interaction (PPI) networks for disease-associated IT neuron modules. Color scale indicates interaction effect size or the degree of change across time points in zQ175 IT neurons relative to WT. Node size indicates degree of PPI interactions and width of the edges between nodes indicates strength of interaction. **b** 6- and 18-month zQ175 vs. WT effect sizes for top hub genes shared between WGCNA and PPI networks. **c** Gene overlap between motor cortex CPN subpopulations in human HD from Pressl et al.^29^ and IT module genes, separated by the direction of effect reported in the human deep layer cortical neurons. **d** Differential TF regulon analysis in zQ175 vs. WT IT neurons at 6- and 18-months. Scatter plots show log_2_ fold change values for negatively correlated genes (x-axis) vs. positively correlated genes (y-axis) of significant TF regulons. Upper left quadrant: repressed TF regulons; Bottom right quadrant: activated TF regulons.

IT-M6 was strongly enriched for proteostasis-related processes. While striatal SPNs showed early proteostasis dysfunction coinciding with huntingtin aggregate emergence prior to 6 months of age (Supplementary Fig. 1), cortical IT neurons exhibited a delayed response. By 18 months key mediators (e.g., *Hspa8, Syt11, Hsp90aa1, Gapdh*) were upregulated, as was the majority of the network (Fig. 7a-b). This delayed timing relative to SPNs may reflect differences in aggregate burden, as cortical huntingtin inclusions appear later and with lower frequency than in the striatum (Supplementary Fig. 1). Together modules IT-M6 and SPN-M5 capture the coordinate decline of protein quality control machinery across regions. We saw a significant overlap of IT-M6 hub genes in L5a human CPNs^29^ and conserved downregulation of hub genes including *SYT11* and *COPG2*, involved in synaptic vesicle release and protein trafficking, suggesting a more widespread breakdown of protein homeostasis and synaptic signaling (Fig. 7). TF regulatory analysis in IT neurons identified increased regulon activity of IT-M6 associated *Neurod2* by 18 months, which was progressively increased in SPNs from 6 to 18 months (Fig. 7d).

IT-M9 shared multiple hub genes with disease-associated striatal modules and human signatures^29^ (e.g., *Pde10a, Phactr1, Kcnd2, Plcb1, Kalrn, Arpp21, Ralyl*), and was progressively downregulated across IT neurons (Fig. 7a-b and Supplementary 8c). The well-characterized biomarker, *Pde10a* showed a significant effect of genotype over time, progressively decreasing, consistent with the progressive dysfunction of IT neurons (Fig. 7b).

## DISCUSSION

This temporal single-cell resource distinguishes stage-specific transcriptional programs across the corticostriatal circuit, revealing how interconnected neuronal populations fail sequentially in HD. Through stage-specific profiling, we distinguish core pathogenic mechanisms from cell type-specific adaptations associated with hierarchical patterns of disease vulnerability.

Consistent with prior observations of early striosomal compartment vulnerability, we demonstrate a dynamic erosion of compartmental identity over time that is distinct and accelerated relative to the canonical D1/D2 vulnerability axis. Loss of the striosomal-matrix distinction has emerged as a core pathological hallmark, accompanied by progressive molecular remodeling within the matrix compartment. Notably, late-stage matrix SPNs downregulated striosomal markers and aberrantly upregulated neurodevelopment- and pluripotency-associated genes, suggesting maladaptive reactivation of developmental programs. These data expand on earlier reports of striatal transcriptional de-repression and epigenetic remodeling in mouse models and human HD^10,95^. To systematically characterize these progressive changes, we applied network-based approaches to identify coherent transcriptional programs.

Using a high-dimensional WGCNA method, we identified cell type-specific gene modules in two vulnerable neuronal populations, SPNs and cortical IT neurons, representing distinct disease-associated transcriptional programs with subtype-specific resolution. The 12 SPN co-expression modules define functionally coherent disease programs, nine of which were preserved in other mouse models and human HD datasets, validating their translational relevance. These modules further resolve the temporal dynamics of previously reported HD transcriptional changes across pathway and compartment SPN identities. Among these, we identify progressive dysregulation of gene networks involved in protein quality control, RNA splicing, GPCR signaling, and synaptic maintenance (SPN-M2, M4, and M6), distinct from divergent age-related responses (SPN-M5 and M7) and late-emerging mechanisms (SPN-M12). We further identified the underlying drivers of dynamic temporal shifts in disease gene programs, including direct mHTT interactor-mediating processes such as the accumulation of nuclear mHTT inclusions (SPN-M7) and failure of early synaptic compensatory responses (SPN-M11). Together, these modules provide a structural framework for understanding the progressive, biphasic, and preterminal phases of SPN degeneration.

Extending our analysis beyond the striatum, our cortical profiling represents the first single-cell cortical analysis of a full-length mHTT model, highlighting disease heterogeneity in both mouse models and human post-mortem HD cortex. Here, we provide a focused analysis of a vulnerable subpopulation of cortical pyramidal neurons that send projections to striatal targets (IT neurons). Our results demonstrate a temporally dynamic shift in BDNF-TrkB signaling not seen at 6 months in the zQ175 mouse model, indicative of the progressive loss of neurotrophic support. Furthermore, gene co-expression analysis within IT neurons identifies layer-specific dysregulation of synaptic signaling, axonal maintenance, and alternative splicing. Cross-module analysis between the striatum and cortex reveal shared disease mechanisms in transient gene signatures (SPN-M7, SPN-M11, IT-M6, and IT-M8), which capture temporally delayed disease processes in cortical neurons involved in proteostasis, glutamatergic signaling, and metabolism. At the gene-level, we delineated region-specific alterations in glutamate receptor composition and potassium channel expression, further resolving the temporal evolution of cortical hyperexcitability, consistent with physiological observations in the motor cortex of the zQ175 mouse model^96^.

Shared hub genes across striatal and cortical networks implicate common regulatory mechanisms in corticostriatal circuit-level failure. Key regulators include Zswim6, whose partial loss in adult mice has been implicated in SPN synaptic deficits and corticostriatal connectivity^75^. Zswim6 can interact directly with PRC2 to direct chromatin remodeling in striatal SPNs^75^ and is downregulated in human HD post-mortem cortex and striatum^29,30^, implicating it as a potential therapeutic target. Beyond epigenetic modifiers, splicing regulators emerge among the most dynamic core hub genes in both regions, including the RNA-binding proteins Celf2, Celf4, and Rbfox1. The splicing regulator Zranb2 was a top hub gene in SPN-M2, enriched for RNA splicing and spliceosome genes, with substantial upregulation in 18-month SPNs. Intriguingly, Zranb2 interacts with Drd2 to influence short- and long-isoform usage^97^, suggesting pathway-specific splicing dysregulation may contribute to D2-MSN vulnerability.

Transcriptional regulatory network analysis revealed stage-specific changes in TF activity during progressive SPN dysfunction. At 6 months, was a decrease in the activity of stress response regulators such as Foxo1 and altered activity of SPN specification factors (Isl1, Wt1), consistent with impaired cellular adaptation and SPN identity loss as an early event in SPN pathogenesis. By 18 months, the predicted activation of neurodevelopmental regulators (Neurod2, Shox2, Bach1) supports reactivation of developmental programs following early dysregulation of SPN specification. Notably, Neurod2 was predicted to repress Foxo1 while activating other developmental TFs, suggesting a feed-forward mechanism that could contribute to the hierarchical breakdown of SPN identity and de-repression of neurodevelopmental transcriptional programs in HD.

These findings reveal the hierarchical, temporally staged transcriptional mechanisms associated with disease progression from early stress adaptation to progressive synaptic dysfunction and loss of neuronal distinctions. We identified conserved compartmental, pathway, and regional dynamics at the gene, module, and regulatory levels in other mouse and human datasets. We have made the results from this study available for exploration in an interactive web portal (see Data availability). This resource establishes a framework for hypothesis-driven functional studies and identifies candidate therapeutic targets for stage-specific interventions in HD.

## METHODS

### Animals

B6J.zQ175 KI mouse lines were obtained from Jackson Laboratories and maintained on a C56B6/J background. Mice were genotyped with primers human *HTT* exon 1 specific primers, and age-matched WT littermates were used as controls for all experiments. All mice were housed in groups of up to five animals per cage and in enriched, temperature-controlled environments with a 12-hour light/ dark cycle. Food and water were provided ad libitum. All animal work adhered to NIH guidelines under IACUC approved by protocols at the Children’s Hospital of Philadelphia.

### HTT CAG Repeat Sizing

DNA from micro dissected cortex and striatum of 6-month and 18-month zQ175 mice was extracted using the Qiagen Puregene cell core kit. The extended HTT repeat was PCR amplified from the purified DNA and sent to Genewiz for capillary electrophoresis (CE). CE data was analyzed, and representative spectra were created using TRACE (https://traceshiny.mgh.harvard.edu/app/traceShiny).

### EM48 Immunohistochemistry

Mice were maintained on a 12 h light/dark cycle under standard temperature and humidity conditions with ad libitum access to food and water. Eighteen- and six-month-old zQ175 mice (n = 3 per age group) were anesthetized with isoflurane and transcardially perfused with 0.05 M PBS (pH 7.4), followed by 50 mL of 4% (w/v) paraformaldehyde in 0.1 M phosphate buffer (pH 7.4). Brains were cryoprotected, coronally sectioned, and processed for immunohistochemistry.

All incubations were performed in PBS containing 0.3% (v/v) Triton X-100 (Triton-PBS). Sections were blocked for 2 h at room temperature (RT) with gentle agitation in Triton-PBS containing 10% (v/v) normal goat serum (NGS), then incubated overnight at RT with anti-EM48 antibody (1:350; Millipore, MAB5374). Following PBS washes, sections were incubated for 4–6[hat RT in goat anti-mouse Alexa Fluor 546 secondary antibody, and counterstained with DAPI. Sections were mounted on glass slides using Fluoromount-G (Southern Biotech), and images were acquired using a Leica SP8 confocal microscope.

### Single nuclei isolation and fixation

All procedures were performed on ice. 75-100 mg micro-dissected tissue samples obtained from murine motor cortex and striatum were incubated in 500 uL of homogenization buffer (320 mM sucrose, 5 mM CaCl_2_, 3 mM Mg(Ac)_2_, 0.1 mM EDTA [pH 8.0], 10 mM Tris HCL [pH 8.0], 10% Triton X-100, 0.2 U/uL SUPERaseIN RNase Inhibitor, 0.1 U/uL RNaseOUT RNase Inhibitor, 0.1 mM Dithiothreitol [DTT]) for 10 min. on ice. Tissue homogenates were then transferred to 2 mL KIMBLE® Dounce All-Glass Tissue Grinders and further homogenized using the loose (A) pestle (10 strokes for striatum, 25 strokes for cortex), followed by 10 strokes using the tight (B) pestle. Homogenates were subsequently passed through a 40-micron cell strainer and collected into pre-chilled 1.5 mL RNase/DNase-free Eppendorf® tubes. Tissue grinders were washed with an additional 500 uL of homogenization buffer which was also passed through a 40-micron cell strainer to improve final nuclei retention. Nuclei were pelleted by centrifugation at 600 x g for 10 min. at 4°C. Supernatants were removed and nuclei pellets were resuspended in wash buffer (1X PBS, 1% BSA, 0.4 U/uL SUPERaseIN RNase Inhibitor, 0.2 U/uL RNaseOUT RNase Inhibitor). The nuclei were pelleted by centrifugation at 600 x g for 5 min. at 4°C, the wash step was repeated twice more, for three washes total. Nuclei pellets were resuspended in 500 uL of wash buffer, passed through a 40-micron Scienceware® Flowmi™ cell strainer, and collected into pre-chilled 1.5 mL RNase/DNase-free Eppendorf® tubes.

Final nuclei suspensions were pelleted by centrifugation at 600 x g for 5 min. at 4°C and resuspending in 750 uL wash buffer. Nuclei quality was assessed by staining an aliquot in DAPI and imaging on EVOS microscope at 10X and 40X magnification. For long-term storage, we adapted a protocol from Rosenberg et al ^35^. Nuclei were fixed in 10 mL 1% formaldehyde in 1X PBS solution for 10 min. then permeabilized with the addition of 40 uL 5% Triton X-100 for 3 min. After centrifugation at 500 x g for 5 min. at 4°C in a swinging bucket rotator and resuspended in 1 mL neutralization buffer (1X PBS, 10 mM Tris-HCL [pH 8.0], 0.2% Triton X-100). A final centrifugation was performed at 500 x g for 5 min. at 4°C in a swinging bucket rotator and nuclei were resuspending in ∼150-300 uL of nuclei storage buffer (1X PBS, 0.2 U/uL SUPERaseIN RNase Inhibitor, 0.1 U/uL RNaseOUT RNase Inhibitor). DAPI+ nuclei were counted via hemocytometer on an EVOS microscope and diluted to ∼2500 nuclei/ uL in a 500 mL cryovial. A final concentration of 5% DMSO was added dropwise with gentle agitation and cryovials were stored in a Corning® CoolCell™ Freezer Container to cool at a rate of - 1°C/minute for 24 hours prior to long-term storage at -80°C. Frozen-fixed nuclei suspensions were stored for up to 6 months.

### SPLiT-seq sublibrary generation

Split-pool ligation-based barcoding was performed as described by Rosenberg et al.^35^, with in-house optimizations at multiple steps. All barcoding steps were performed in 96-well DNase/RNase-free low-bind polypropylene plates. The first round of barcoding added a well-specific barcode by an *in situ* reverse transcription (RT) reaction with 48 well-specific barcoded poly(dT)15 RT primers. Nuclei were diluted to 750 nuclei/uL with 1X PBS + RNase Inhibitor ( 0.2 U/uL SUPERaseIN RNase Inhibitor, 0.1 U/uL RNaseOUT RNase Inhibitor) and loaded at a volume of 8 uL for a final loading of 6,000 nuclei per well into an RT mix (4 uL 5X RT Buffer, 0.625 uL of RNase-free water, 1 uL of 40mM dNTP mix, 4 uL of 25 μM poly(dT)15 RT primer, 2 uL Maxima H Minus Reverse Transcriptase). The Round 1 plate was incubated in a thermocycler for 10 min at 50°C before cycling three times at 8°C for 12s, 15°C for 45s, 20°C for 45s, 30°C for 30s, 42°C for 2 min, and 50°C for 3 min, followed by a final step at 50°C for 5 min. The plate was placed on ice, and all RT reactions were pooled into a pre-chilled 15 mL falcon tube. 9.6 μL of 10% Triton X-100 was added to the nuclei suspension before centrifugation at 500 x g for 10 min. at 4°C. The supernatant was discarded, and nuclei were resuspended in 2 mL 1X NEB buffer 3.1 and 20 uL of RNaseOUT RNase Inhibitor.

Round 2 and 3 barcoding plates were prepared according to the original protocol. The nuclei suspension was added to a ligation mix (500 uL 10X T4 Ligase buffer, 100 uL T4 DNA Ligase, 40 uL RNaseOUT RNase Inhibitor, 20 uL SUPERaseIN RNase Inhibitor, 100 uL 5% Triton-X 100, up to 2 mL RNase-free water) and distributed onto the Round 2 barcoding plate (40 uL per well) with 26.4 μM 10 uL of well-specific DNA barcodes. The ligation reactions were incubated at 37°C in a thermocycler for 30 min. 10 uL of blocking oligonucleotide (BC_0126) in 2.5X T4 Ligase Buffer was added to each well and incubated for 30 min. at 37°C to prevent further ligation of unbound DNA barcodes. Nuclei were pooled together from all wells and passed through a 40-micron cell strainer into a reagent reservoir. 100 uL of T4 DNA Ligase was added to the pooled nuclei suspension before distribution onto the Round 3 barcoding plate with well-specific DNA barcodes containing unique molecular identifiers (UMIs), a universal PCR handle, and a biotin molecule for future purification (60 uL final volume per well). After a final 30 min. incubation at 37°C, blocking oligo (11.5 μM, BC_0066) was added with 125 mM EDTA in RNase-free H_2_O to terminate the reaction. Nuclei were pooled together into a 15 mL conical tube and 70 uL of 10% Triton-X 100 was added prior to centrifugation at 1000 x g for 10 min. at 4°C. The nuclei pellet was resuspended in a wash buffer (4 mL 1X PBS, 40 uL 10% Triton-X 100, 10 uL SUPERaseIN RNase Inhibitor) and centrifuged again at 1000 x g for 5 min. at 4°C.

The nuclei pellet was resuspended in 50 uL 1X PBS + RNase Inhibitors and a 5 uL aliquot was taken for dilution in 1 ug/mL DAPI in 1X PBS for final nuclei counting via hemocytometer. Nuclei were diluted to 5,000 nuclei (25 uL 1X PBS + RNase Inhibitors) in PCR 8-tube strips. 25 uL of 2X lysis buffer (10 mM Tris HCL [pH 8.0], 5 M NaCl, 0.5 M EDTA [pH 8.0], 10% SDS, RNase-free H_2_O) and 5 uL of 20 mg/ml Proteinase K was added to each sublibrary. Final nuclei sub-libraries were lysed via a 120 min. incubation at 55°C in a thermocycler and stored at -80°C for up to 3 months.

### cDNA sublibrary template-switching and amplification

Nuclei sub-libraries were thawed in a 37°C bead bath and then incubated for 60 min. at room temperature with constant agitation (∼ 600 RPM) in Streptavidin MyOne C1 beads (according to manufacturers’ protocol) to purify Biotin-tagged cDNA. A template-switching reaction to add a 5’ universal PCR handle was performed in an RT reaction mix (44 uL RNase-free H_2_O, 22 uL Maxima RT Buffer, 22 uL 20% Ficoll PM-400, 11 uL 10 mM dNTP mix, 2.25 uL RNaseOUT RNase Inhibitor, 2.25 Template-Switching Oligo (TSO; BC_0127), 5.5 uL Maxima RT H Minus Enzyme). Sub-libraries in TSO mix were incubated at room temperature for 30 min. after which the beads were resuspended, and the reactions were incubated for 90 min. at 42°C in a thermocycler. cDNA-bound beads were washed with 125 mL of 10 mM Tris-HCL, 1% Tween-20, and SUPERaseIN RNase Inhibitor. cDNA sub-libraries were amplified by polymerase chain reaction (PCR) with primers BC_0108 and BC_0062 in KAPA HiFi 2X master mix. Initial denaturation at 95°C for 3 minutes, was followed by five cycles at 98°C for 20 seconds, 65°C for 45 seconds, and 72°C for 3 minutes. This was succeeded by a second round of qPCR with thermocycling conditions of 95°C for 3 minutes, followed by cycling at 98°C for 20 seconds, 65°C for 20 seconds, and 72°C for 3 minutes, with a final extension at 72°C for 5 minutes. A 0.8X SPRI bead clean-up was performed post-PCR amplification, and sub-libraries were stored at ∼10-50 ng/uL in RNase-free H_2_O at -20°C overnight. Amplified cDNA sublibrary concentration was measured by Qubit and size distribution with an Agilent D5000 tape station.

### cDNA sublibrary fragmentation and library indexing

The protocol for cDNA fragmentation, end-repair, A-tailing, and Illumina indexing was adapted from the manufacturer’s manual for the KAPA HyperPlus kit. 100 ng of cDNA was diluted in 50 uL of fragmentation buffer (10 mM Tris-HCL [pH 8.5], 1X KAPA Fragmentation Buffer and Enzyme) and incubated for 10 min at 32°C in a thermocycler. 1X ER & A-tailing Buffer and Enzyme mix were added and the reactions were incubated at 65°C for 30 min. A ligation mix was added to each reaction containing Illumina adapters (50 uM Illumina adapter stock (BC_0243 and BC_0244), KAPA ligation buffer, KAPA DNA ligase, PCR-grade H_2_O) and incubated for 15 min. at 20°C. A 0.8X SPRI bead cleanup was performed, and cDNA was eluted in 21 uL of RNase-free H_2_O. Sublibrary-specific indexed PCR primers (BC_0070 – BC_0093) and universal index (BC_0027) were added with 2X KAPA HiFi Master Mix. The PCR reaction was as follows, an initial denaturation at 95°C for 3 minutes, followed by ten cycles at 98°C for 20 seconds, 67°C for 20 seconds, and 72°C for 3 minutes, and a final extension at 72°C for 5 min. A two-sided SPRI bead cleanup was performed to select for cDNA between 300-600 bp (0.6 – 0.8X).

### Illumina Sequencing

Libraries were sequenced on an S2 NovaSeq v1.5 6000 flowcell using 200-bp paired-end sequencing kit. Our desired sequencing depth of 50-60 thousand reads per nuclei was determined by first deep sequencing a sublibrary of 500 nuclei at a total depth of 200,000 reads per nuclei and sub-sampling the raw FASTQ files to determine the point of sequencing saturation. Read 1 (106 bp) was used to map transcript sequences, while Read 2 (94) contained the single-cell barcode and UMI for read demultiplexing.

### SPLiT-seq single-nucleus read demultiplexing

To process SPLiT-Seq barcoded single-cell datasets we developed SPLiT-Seq_Demultiplexing, an open-source, portable, and high-throughput pipeline for end-to-end assignment of SPLiT-Seq barcodes. This tool enables the conversion of raw barcoded FASTQ files to gene x cell count matrices, the most common input into downstream single-cell analysis programs. Here, the python-based version of SPLiT-Seq demultiplexing (splitseqdemultiplex_0.2.3.py) was run once for each Illumina indexed sublibrary. Each sublibrary was processed using the following optional parameters:

-n 20: The “-n” flag indicates the number of threads used for parallelization of the pipeline. To facilitate parallelization SPLiT-Seq_Demultiplexing splits the input FASTQ file into the provided number of segments. The barcode extraction process is then performed in parallel on all segmented FASTQ files.

-e 1: The “-e” flag passes the permissible error rate for demultiplexing. Here an error rate of 1 was used meaning that each 8bp (Round 1, 2, or 3 barcode) must be within an edit distance of 1 from a known SPLiT-Seq barcode sequence. If a barcode contains an error but passes this filter it is corrected to the sequence of the known barcode.

-m 200: The “-m” flag indicates the minimum number of reads required per barcode. 200 is a recommended threshold when processing production scale datasets as higher thresholds improve performance and greatly reduce output count matrix size by limiting the number of low diversity cells carried forward.

Other input selections are required but have less direct impact on output and are described in the package documentation.

Using the above parameters SPLiT-Seq_demultiplexing performs parallelized, position-based barcode extraction by first automatically learning the position of the Round 1-3 barcodes from read2 of the Illumina paired-end sequencing output using static barcode flanking sequences. If a complete set of SPLiT-Seq barcodes 1-3 along with their associated UMI are extracted from Read 2 they are then appended to the ID of the Read 1 mate-pair, which contains transcriptomic information. If no complete set of barcodes and UMI are found, then the read is discarded. The above extraction process is performed and written to disc in user-defined bins to control memory usage. The output of all passing reads being processed across separate threads is merged into a single output. A FASTQ file containing the read information from Illumina Read 1 and the barcode information extracted from read 2 appended to the Read1 read ID is outputted.

Performance metrics are calculated from this single FASTQ file and minimum read filtering is applied. Mapping is then performed with STAR (2.6.0), feature assignment is performed with FeatureCounts (2.0.3), format conversion is performed using samtools (1.9), and UMI collapse is performed with UMI-tools (0.5.5).

### Single-nuclei RNA-seq quality control and clustering

Individual raw count matrices corresponding to each sublibrary were imported into R (v4.3.1) and individual Seurat (v5.1.0) objects were constructed. Sample-level metadata was integrated with each Seurat object based on the 8-bp Round 1 barcode, excluding nuclei with less than 500 UMIs detected, and genes identified in fewer than 10 nuclei. All sublibrary-level Seurat objects were merged for downstream analysis, then subset by region. Nuclei with greater than 5% of reads mapping to mitochondrial genes or greater than 15000 UMIs detected were excluded from further analysis, yielding 26,283 cortical and 18,861 striatal high-quality nuclei.

The gene expression matrix was log normalized and scaled in Seurat (v5.1.0). The top 2000 most variable genes were identified using variance-stabilizing transformation, excluding mitochondrial genes. Principal Component Analysis (PCA) was used to reduce the dimensionality of the scaled gene expression matrix, after regression of UMI count and mitochondrial percentage. Samples were integrated using Harmony (v1.2.3), first by biological replicate for annotation then separately by sublibrary for visualization and downstream analysis to retain variance by experimental condition. We chose to harmonize by replicate for clustering and annotation to avoid cell type classification of clustering due to genotype or age. The first 20-30 harmony reductions were used to construct a K-nearest neighbor (KNN) graph, followed by iterative, Louvain clustering. Doublets were identified using scDlbFinder (v1.18.0) with a predicted doublet rate of 7.5% and sample-specific detection. Only singlets were retained for downstream analysis. UMAP visualized used Harmony dimensions (cortex: dimensions = 25, n. neighbors = 30, min. dist = 0.4, metric = cosine; striatum: dimensions = 20, n. neighbors = 20, min. dist = 0.4, metric = cosine).

### Reference-based cell type mapping and annotation

Clusters were annotated using reference-based mapping and validated by canonical marker gene expression. Cortical nuclei were mapped using Azimuth^32^ (v.0.5.0) and the Azimuth mouse motor cortex reference dataset ^32^, and secondarily using Hierarchical nearest neighbor (HANN) mapping to the 10X whole mouse brain taxonomy ^98^ using the Allen Brain Institute “MapMyCells” web application (RRID:SCR_024672). The same tool was used to annotate striatal nuclei, mapping nuclear profiles to the Human and Mammalian Brain Atlas (HMBA) basal ganglia consensus taxonomy. Annotations were confirmed by region-specific cell type-specific marker genes. Low quality clusters (low UMI or high mitochondrial content), ambiguous populations, and putative doublets were excluded through iterative clustering and quality control analysis.

### Cell marker gene identification

We performed a differential gene expression analysis comparing the gene expression of each cell type against the remainder, or a one-versus-all approach, to identify marker genes with the Seurat function FindAllMarkers. We chose to use a single-cell hurdle model, MAST^99^, accounting for sublibrary batch, sex and UMIs per cell as a covariate. We filtered our results to only include protein-coding genes expressed in at least 10% of nuclei with a log_2_ fold change ≥ 0.25 and Bonferroni adjusted *p*-value < 0.001. When establishing marker gene expression for comparison in zQ175, only WT 6-month nuclei were used for analysis.

### Differential cell type abundance analysis

To assess changes in cell type proportions experimental conditions, we performed differential abundance testing using EdgeR^47^ (v4.1.2). We performed this analysis separately for motor cortex and striatum at both time points (6 and 18 months). fitted a Negative Binomial Generalized Linear Model (NB GLM) for count data (nuclei per annotation) using the edgeR^68^ package (v4.0.16). Biological replicates with less than 250 nuclei were filtered prior to analysis of cell composition. Our design matrix was as follows ∼ Sublibrary + Sex + Condition, accounting for technical variation in library preparation. Differential abundance was assessed by the glmQLFTest function. We adjusted for multiple comparisons with the Benjamini-Hochberg procedure and set our significance threshold at a false discovery rate (FDR) < 0.05.

### Pseudocell differential gene expression analysis

To increase statistical power for differential gene expression analysis while preserving biological variability, we utilized a pseudocell approach^49,50^ accounting for technical variability and dropout rate in sparse single-cell data. We adapted a published pipeline^49,50^ to include analysis of age-genotype interactions.

“Pseudocells” were constructed by grouping single nuclei transcriptomes by replicate, cell type, and brain region and aggregating raw expression matrices using a k-means clustering of the top 30 principal components. For cell type-specific analyses, pseudocells were generated every 25 nuclei (10 nuclei minimum), whereas for cell class-level analyses pseudocells were generated every 50 nuclei (25 nuclei minimum). Cell types were excluded with less than 20 nuclei per cell type per replicate. Replicates were excluded with less than 2 pseudocells.

Differential gene expression analysis was performed using a linear mixed-effects model framework implemented through the limma-voom pipeline with sample quality weights. Sex, SPLiT-seq library batch, and pseudocell size were accounted for as covariates and replicate ID as a random effect to account for individual variance. The model included a 4-level group factor representing all combinations of age (6 months, 18 months) and genotype (WT, zQ175).

Four contrasts were tested using empirical Bayes moderated t-statistic: (1) zQ175 vs. WT at 6 months, (2) zQ175 vs. WT at 18 months, (3) aging effect in WT mice (18 months vs. 6 months), and (4) age-genotype interaction (zQ175 18 months – zQ175 6 months) – (WT 18 months – WT 6 months). The interaction term identifies genes with progressive or stage-specific disease effects that differ from normal aging. P-values were adjusted for multiple testing using the Benjamini-Hochberg false discovery rate method (adj. *p* < 0.05, |log_2_FC| < 0.1). Significance for interaction effects was defined as adj. *p* < 0.05 and |log_2_FC| < 0.2.

To classify robust temporal gene expression signatures, we added additional criteria:

1. “Directional Opposition” - genes with opposing genotype effects at both timepoints (|log_2_FC| ≥ 0.25 at each timepoint with sign reversal);
2. “Divergent” - genes non-significant at 6 months (adj.p > 0.05) but significantly dysregulated at 18 months (adj.p < 0.05);
3. “Convergent” - genes with early significant dysregulation (6 months adj.p < 0.05) that normalized by 18 months (adj.p > 0.05).

All patterns except Directional Opposition required consistent directionality across timepoints.

### Pseudobulk differential gene expression analysis

To validate the robustness of our pseudocell-based approach for differential expression testing, we performed parallel pseudobulk analysis. For each cell type and biological replicate, all raw expression matrices were aggregated into a single pseudobulk sample using the aggregateAcrossCells function from the scuttle package. Genes were filtered to a minimum of 10 counts to remove lowly expressed genes. We utilized the same approach via limma-voom with sample quality weights, but unlike the pseudocell model, replicates were not accounted for as a random effect as each mouse contributes a single pseudobulk sample per cell type. Library batch and sex were included as covariates.

Concordance between pseudocell and pseudobulk approaches was assessed by calculating Pearson correlation for log_2_ fold changes and Spearman correlation for *p*-values across all contrasts and cell types (Supplementary Dataset 2).

### Gene Set Enrichment Analysis

Gene set enrichment analysis was performed using the R package FGSEA. DEGs were filtered to include only protein coding genes and ranked by t. statistic for each comparison described.

### Jensen-Shannon Divergence (JSD) Analysis

To quantify transcriptomic distance between cell populations, we calculated the Jensen-Shannon distance as adapted from Matsushima et al.^20^. To reduce noise, we defined the average gene expression for all genes expressed in ≥ 5% of nuclei in each condition. Nuclei profiles were subsampled to maintain equal numbers across samples and conditions within each comparison. The philoentropy R package was used to calculate the pairwise JSD between SPN subtypes. The difference in JSD was defined the distance between zQ175 and WT and was used to quantify transcriptional identity loss in Fig. 3. Negative values indicate reduced distinction between a given SPN axis in zQ175 mice.

### CAG repeat-length dependent gene signatures

The expression of CAG repeat-length dependent gene signatures described in Handsaker et al.^10^ were calculated using the R package UCell (v 2.8.0), utilizing a Mann-Whitney U statistic approach to evaluate the enrichment of a gene set in individual cells, then ranked by expression levels and compared against a background set of randomly selected genes. This method is robust to differences in library size. The AddModuleScore_UCell wrapper function for Seurat objects was used to score nuclei for expression of positive correlated, CAG (+), and negatively correlated, CAG (-), gene sets in a given dataset. Scores range from 0 to 1, higher values indicating greater enrichment of that gene signature in a given cell.

### hdWGCNA co-expression network analysis

The hdWGCNA ^73^ (v0.4.08) R package was used to carry out weighted gene co-expression network on cell type-specific subpopulations: SPNs and IT neurons. Analysis was restricted to protein-coding genes expressed in at least 5% of nuclei in at least one condition. To account for the sparsity of single nuclei data, we computed “metacells” by aggregating cells from the same biological origin (e.g., mouse, cell type and region). Metacell parameters were optimized for each cell type. For SPNs k = 50 nearest neighbors, requiring a minimum of 100 cells per grouping and allowing a maximum of 20 shared nuclei per metacell. For IT neurons k= 35, requiring 250 cells per grouping and a maximum of 5 shared nuclei per metacell.

A soft-power threshold search was performed to identify the optimal soft power threshold (SPNs, soft power = 9; CPNs, soft power = 6) prior to computing co-expression network topological overlap matrices (TOMs) for each cell type. The ConstructNetwork function was used to identify subsequent gene modules. SPNs: deepSplit = 4 and mergeCutHeight = 0.2; IT neurons: deepSplit = 3 and mergeCutHeight = 0.15. Both analyses used minModuleSize = 40 and networkType = “signed”.

Module eigengenes (MEs), representing the first principal component of each modules’ expression, were calculated using the ModuleEigengenes function. Eigengene-based connectivity (kME) values for each module-associated gene were determined by the hdWGCNA function ModuleConnectivity. Differential module eigengene analysis was performed with Wilcoxon rank sum accounting for IT neurons across pseudotime, separated into 10 bins (Fig. 6). Module-trait correlation with genotype, age, CAG (+) and CAG (-) scores were reported with Pearson correlation coefficients.

Module preservation of SPN modules with SPNs from Lee et al.^30^ (human HD) and Lim et al.^73^ (R6/2) were assessed using the modulePreservation function with 200 permutations. For human datasets, mouse-human gene orthologs were mapped with Ensembl BioMart.

Co-expression networks were integrated with protein-protein interaction data from STRING (v12.0, confidence ≥ 200) by element-wise multiplication of the TOM with binarized PPI adjacency matrices to identify convergent hub genes. Functional enrichment analysis was performed using enrichR (v3.0) on the top 100 hub genes per module against GO Biological Process 2025, GO Molecular Function 2025, KEGG 2021 Human, WikiPathways 2024 Human, and HDSigDB Human 2021 databases.

### Gene overlap analysis

All overlap analyses in this manuscript comparing sets of genes were performed using the R package Gene Overlap (version 1.40.0), calculating the overlap between two gene sets and reporting results from Fisher’s exact test to determine both *p*-values and odds ratios in comparison to a background set of genes. All overlap analyses using the total genes included in the WGCNA analysis as background.

### Transcription regulatory network and regulon analysis

Transcription factor (TF) regulon analysis was performing using hdWGCNA (v 0.2.29). This analysis consists of five main steps: 1) Establish metacells from single-cell data which was done to identify hdWGCNA modules, 2) scan gene promoters for TF motifs, 3) model the gene expression of each predicted target gene as a function of the TF expression, 4) define TF regulons and networks, 5) perform differential regulon analysis to identify perturbed TF regulons across conditions.

TF binding motifs from JASPAR2024 vertebrate CORE collection were scanned across the mouse mm10 genome using motifmatchr with EnsDb.Mmusculus. v79 annotations. TF regulatory networks were constructed using gradient boosting (XGBoost v1.7.7) with parameters: objective=’reg: squarederror’, max_depth=1, eta=0.1, alpha=0.5. Regulons were defined first by a regulatory score of 0.01 and identifying the top 200 target genes per TF. This stringent regulon assignment was used for head-to-head comparisons between TF regulons by differential regulon analysis with Wilcoxon rank sum to identify top targets. Regulons with < 50 genes were excluded from differential regulon analysis. We report also full predicted TF regulons (regulatory score = 0.005) where the top 5 TFs per gene were assigned. For each cell, regulon activity scores were calculated separately for positively correlated target genes (Pearson r > 0.1) and negatively correlated targets (r < -0.1) using the UCell algorithm. These scores represent aggregate target gene expression, providing functional readout of TF activity. Differential regulon activity between zQ175 and WT was assessed at 6- and 18-months using Wilcoxon rank sum tests with Bonferroni correction. TFs were considered differentially active if |log2FC|>0.1 and adjusted p.value was < 0.001 for both positive and negative regulon components.

TF regulatory influence across co-expression modules was quantified by assigning target genes to modules and calculating regulatory relationships as the signed product of regulatory gain and correlation (Gain x sign (Cor). Gain, meaning the importance of a given TFs expression in predicting the expression of a given gene. Only relationships with |regulatory score| > 0.1 were retained. For each TF-module pair, we computed positive and negative links, as well as the regulatory balance (positive – negative counts).

Regulon activity differences across SPN subtypes were tested using linear modules: UCell score ∼ library batch + sex + genotype (within timepoints).

## DATA AVAILABILITY

All raw sequencing data generated as a part of this study will be deposited to NCBI upon publication. All processed count matrices, supplementary data and expanded results tables will be made available.

Publicly available datasets re-analyzed in this study can be found on NCBI GEO query. The Human HD Caudate and Putamen snRNA-seq from Lee et al.^30^ is available under GEO accession: GSE152058, and the R6/2 snRNA-seq study from Lim et al.^73^ in striatum is available under GEO accession: GSE180294.

## CODE AVAILABILITY

All code associated with the SPLiT-seq demultiplexing package will be made available upon publication.

All data processing and analysis code for this study will be made available upon publication.

## ACKNOWLEDGEMENTS

The authors thank Dr. Teodora Orendovici and the rest of the high-throughput sequencing core at CHOP. This study was supported by postdoctoral research fellowships funded by the Hereditary Disease Foundation awarded to P.T.R. and S.S.C., an NHGRI T32 HG000046 (A.B.R.), and an NINDS F31 Ruth L. Kirschstein National Research Service Award F31 NS122297 (A.B.R) and the CHOP Research Institute.

## AUTHOR CONTRIBUTIONS

Conceptualization, A.B.R, P.T.R, B.L.D.; Methodology, A.B.R., P.T.R., I.H., M.K.; Resources, A.B.R.; Software, P.T.R., A.B.R.; Validation, A.B.R.; Investigation, A.B.R., I.H., M.K.; Formal Analysis, A.B.R.; Writing – Original Draft, A.B.R.; Writing – Review & Editing, B.L.D., A.B.R., I.H.; Supervision, B.L.D.

## COMPETING INTERESTS

B.L.D is a founder of Latus Bio; and has sponsored research or consults for Carbon Therapeutics, Latus Bio, and Seamless Ther; P.R.R. is a founder and employee of Latus Bio. The remaining authors declare no competing interests.

## Notes

### Competing Interest Statement

The authors have declared no competing interest.

