## Supplemental Figures for "Temporal single-cell atlas of full-length Huntington’s disease mouse model defines stage-specific signatures of corticostriatal dysfunction"

**Supplementary Information for Robbins et al.**

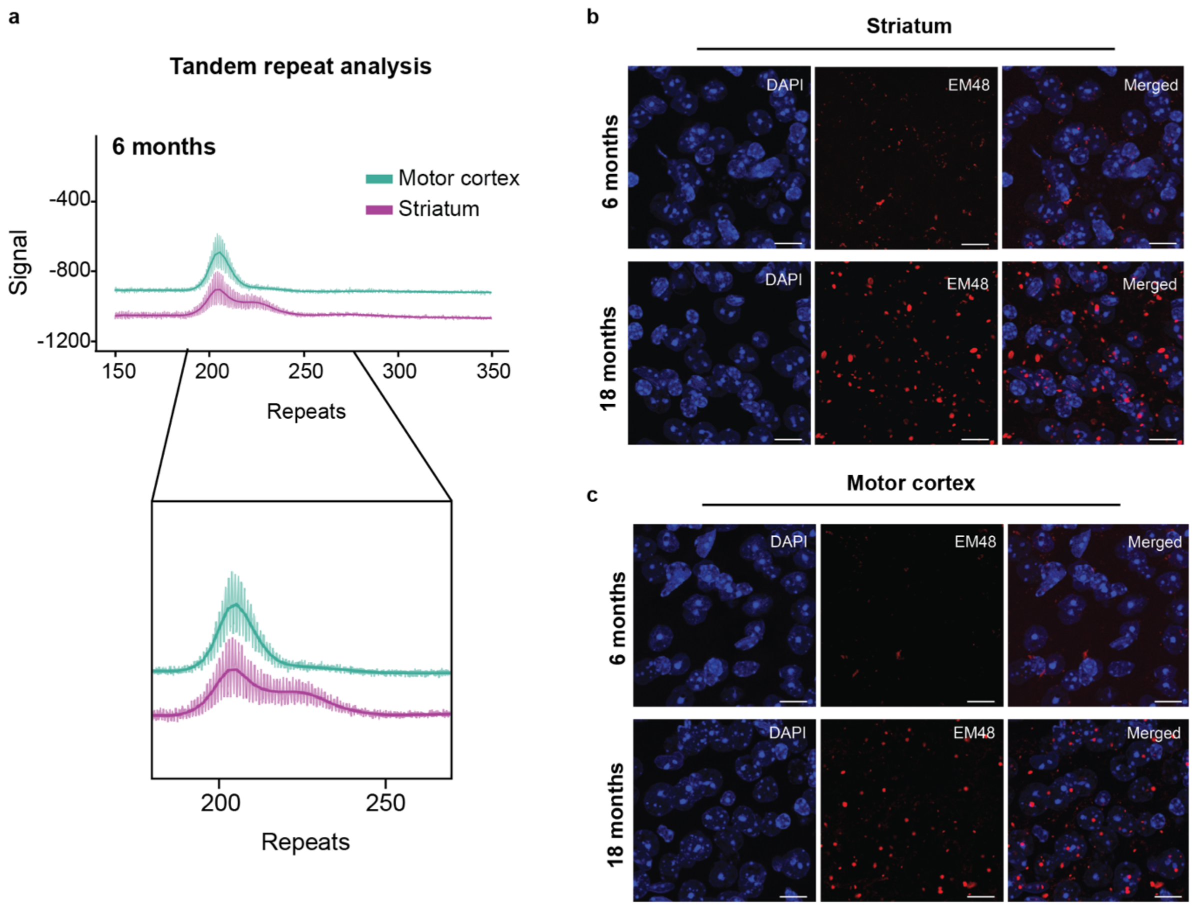


Supplementary Figure 1**. mHTT CAG repeat expansion and aggregation in zQ175 striatum and motor cortex. a** CAG repeat expansion in motor cortex (teal) and striatum (purple) at 6 months of age in zQ175 mice. Bottom panel: magnified view of repeat size distribution peaks. **b-c** Confocal imaging of mHTT inclusions by EM48 staining and DAPI nuclear counterstain in **b** striatum and **c** cortex at 6 and 18 months of age, n = 1 mouse per timepoint. Scale bars, 10 μm.

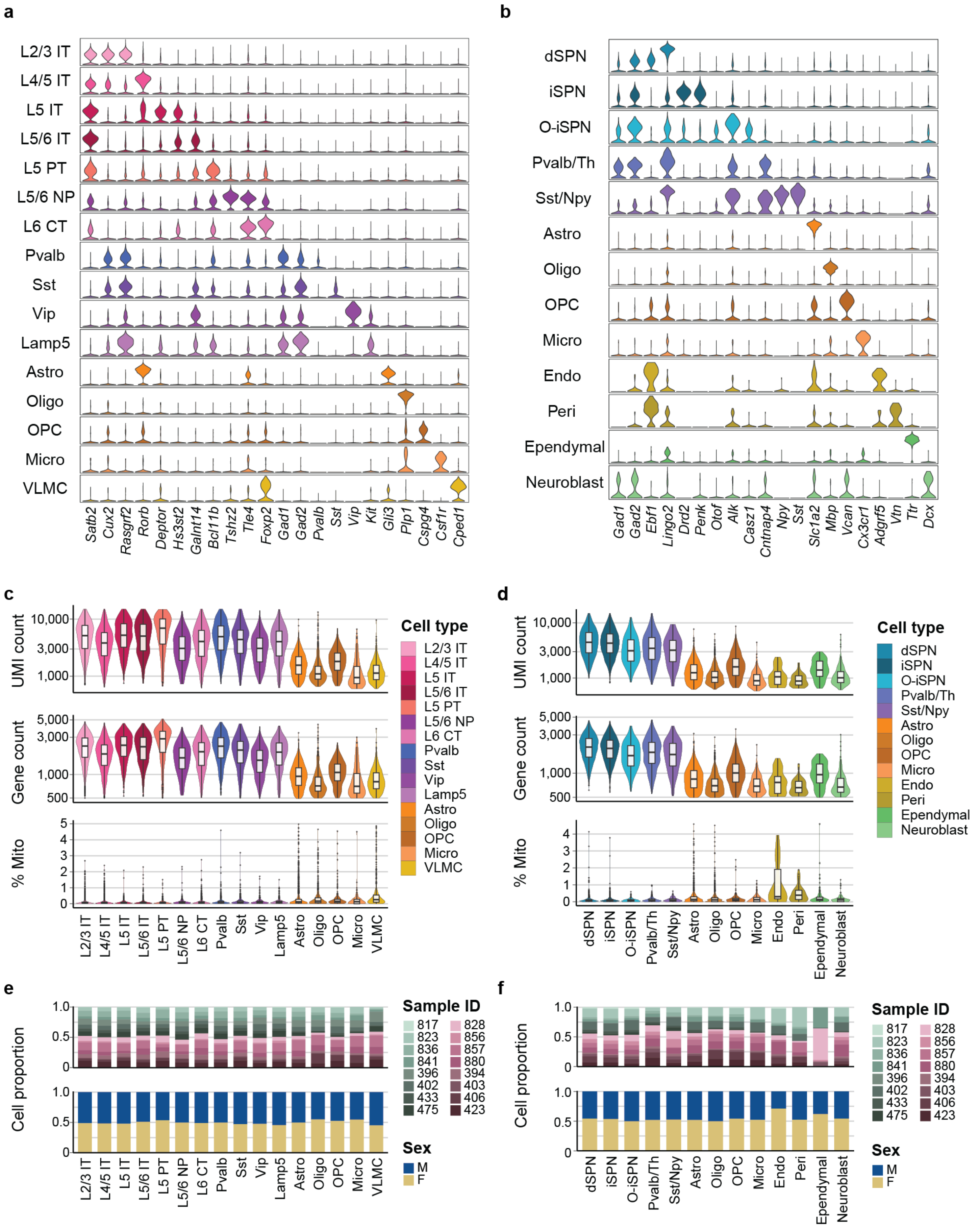

a
L2/3 IT
L4/5 IT
L5 IT
L5/6 IT
L5 PT
L5/6 NP
L6 CT
Pvalb
Sst
Vip
Lamp5
Astro
Oligo
OPC
Micro
VLMC
Satb2 Cux2 Rasgrf2 Rorb Deptor Hs3st2 Galnt14 Bcl11b Tshz2 Tle4 Foxp2 Gad1 Gad2 Pvalb Sst Vip Kit Gli3 Plp1 Cspg4 Csf1r Cped1
b
dSPN
iSPN
O-iSPN
Pvalb/Th
Sst/Npy
Astro
Oligo
OPC
Micro
Endo
Peri
Ependymal
Neuroblast
Gad1 Gad2 Ebf1 Lingo2 Drd2 Penk Otof Alk Casz1 Cntnap4 Npy Sst Slc1a2 Mbp Vcan Cx3cr1 Adgrf5 Vtn Ttr Dcx
c
UMI count
Gene count
% Mito
Cell type
d
e
Cell proportion
Sample ID
817 823 836 841 396 402 433 475
828 856 857 880 394 403 406 423
Sex
M
F
f

Supplementary Figure 2**. Quality control metrics and cell annotation of cortical and striatal samples. a-b** Canonical marker gene expression across cortical (**a**) and striatal (**b**) cell types. Violin plots display the distribution of normalized expression values for each marker across annotated cell types. **c-d** Quality control metrics for cortical (**c**) and striatal (**d**) cell types. Violin plots show distributions of UMI counts, gene counts, and mitochondrial gene percentage for each cell type. Box plots within violins indicate the median (center line), interquartile range(box), containing the middle 50% of the data, and the maximum range extending to 1.5 times the interquartile range. **e-f** Cellular composition by sample and biological sex for cortical (**e**) and striatal (**f**) cell types. Stacked bar plots show the proportion of cells originating from each sample and by biological sex within each cell type.

Abbreviations: IT, Intertelencephalic; PT, pyramidal tract; NP, near-projecting; CT, corticothalamic; Pvalb, parvalbumin; Sst/Npy, somatostatin/neuropeptide Y; Vip, vasoactive intestinal peptide; Lamp5, lysosomal associated membrane protein 5; dSPN, direct spiny projection neurons; iSPN, indirect spiny projection neurons; Astro, astrocytes; Oligo, oligodendrocytes; OPC, oligodendrocyte precursor cells; Micro, microglia; VLMC, vascular leptomeningeal cells; Endo, endothelial; Peri, pericytes.

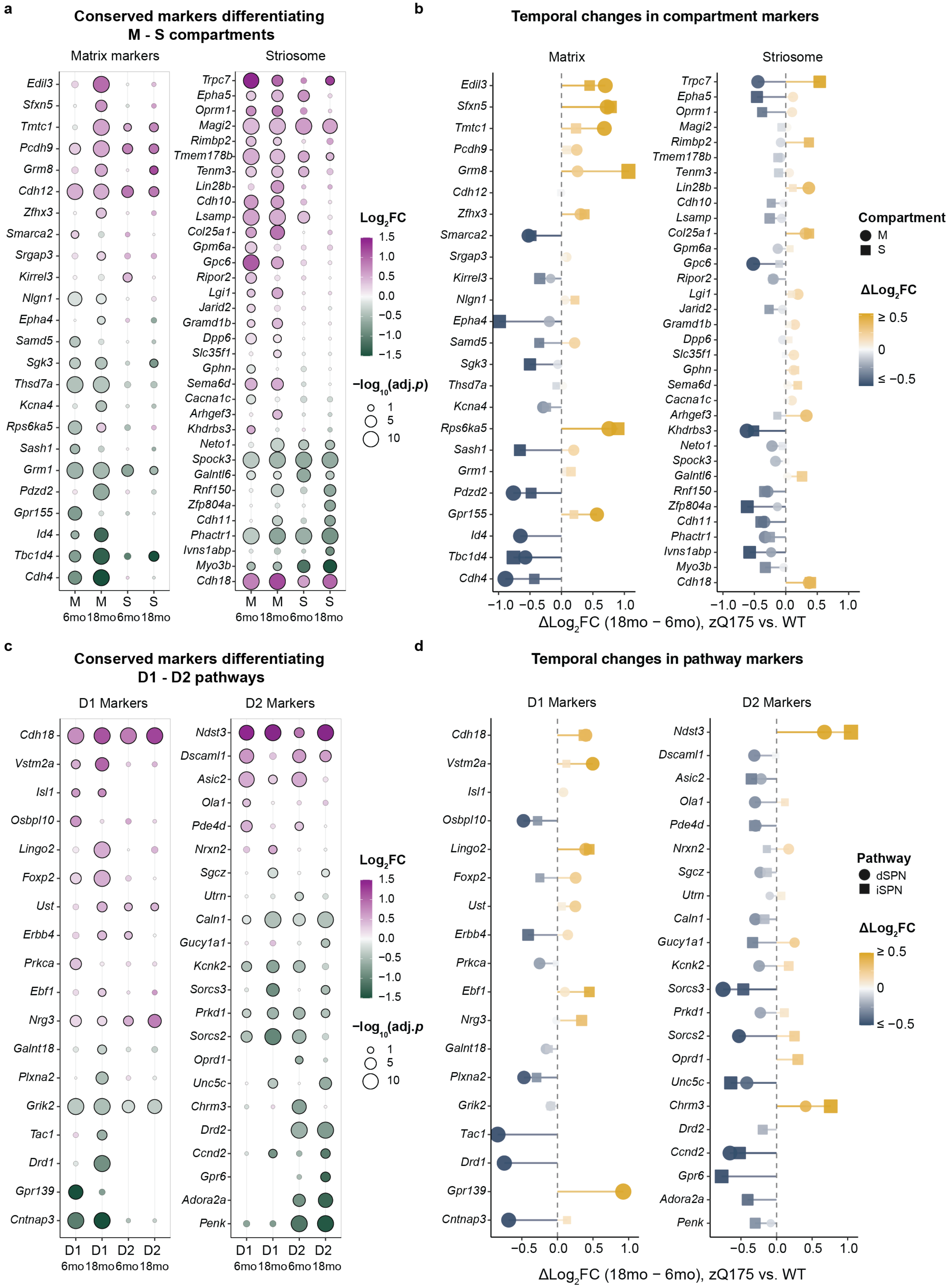
a
Conserved markers differentiating
M - S compartments
Matrix markers
Striosome
Edil3
Sfxn5
Tmtc1
Pcdh9
Grm8
Cdh12
Zfhx3
Smarca2
Srgap3
Kirrel3
Nlgn1
Epha4
Samd5
Sgk3
Thsd7a
Kcna4
Rps6ka5
Sash1
Grm1
Pdzd2
Gpr155
Id4
Tbc1d4
Cdh4
Trpc7
Epha5
Oprm1
Magi2
Rimbp2
Tmem178b
Tenm3
Lin28b
Cdh10
Lsamp
Col25a1
Gpm6a
Gpc6
Ripor2
Lgi1
Jarid2
Gramd1b
Dpp6
Slc35f1
Gphn
Sema6d
Cacna1c
Arhgef3
Khdrbs3
Neto1
Spock3
Galntl6
Rnf150
Zfp804a
Cdh11
Phactr1
Ivns1abp
Myo3b
Cdh18
M M S S
6mo 18mo 6mo 18mo
Log2FC
1.5
1.0
0.5
0.0
−0.5
−1.0
−1.5
−log10(adj.p)
1
5
10
b
Temporal changes in compartment markers
Matrix
Striosome
−1.0 −0.5 0.0 0.5 1.0
ΔLog2FC (18mo − 6mo), zQ175 vs. WT
Compartment
M
S
ΔLog2FC
≥ 0.5
0
≤ −0.5
c
Conserved markers differentiating
D1 - D2 pathways
D1 Markers
D2 Markers
Cdh18
Vstm2a
Isl1
Osbpl10
Lingo2
Foxp2
Ust
Erbb4
Prkca
Ebf1
Nrg3
Galnt18
Plxna2
Grik2
Tac1
Drd1
Gpr139
Cntnap3
Ndst3
Dscaml1
Asic2
Ola1
Pde4d
Nrxn2
Sgcz
Utrn
Caln1
Gucy1a1
Kcnk2
Sorcs3
Prkd1
Sorcs2
Oprd1
Unc5c
Chrm3
Drd2
Ccnd2
Gpr6
Adora2a
Penk
D1 D1 D2 D2
6mo 18mo 6mo 18mo
Log2FC
1.5
1.0
0.5
0.0
−0.5
−1.0
−1.5
−log10(adj.p
1
5
10
d
Temporal changes in pathway markers
D1 Markers
D2 Markers
−0.5 0.0 0.5 1.0
ΔLog2FC (18mo − 6mo), zQ175 vs. WT
Pathway
dSPN
iSPN
ΔLog2FC
≥ 0.5
0
≤ −0.5

Supplementary Figure 3**. Differential gene expression of human HD conserved markers of striatal identity. a** Dysregulation of cross-species conserved matrix (left) and striosomal (right) identity gene markers, significant in at least one timepoint. Significance is denoted by a black outline ($|log_2FC| \geq 0.1$; $p.adj < 0.05$). **b** Temporal changes in compartment gene marker expression. Lollipop plot shows the difference in $log_2FC$ between 18- and 6-month timepoints for the gene markers in (**a**). **c** Dysregulation of cross-species conserved D1 (left) and D2 (right) identity gene markers, significant in at least one timepoint, as in (**a**). **d** Temporal changes in pathway marker expression in (**c**). Lollipop plot shows the difference in $log_2FC$ between 18- and 6-month timepoints for the gene markers in (**c**).

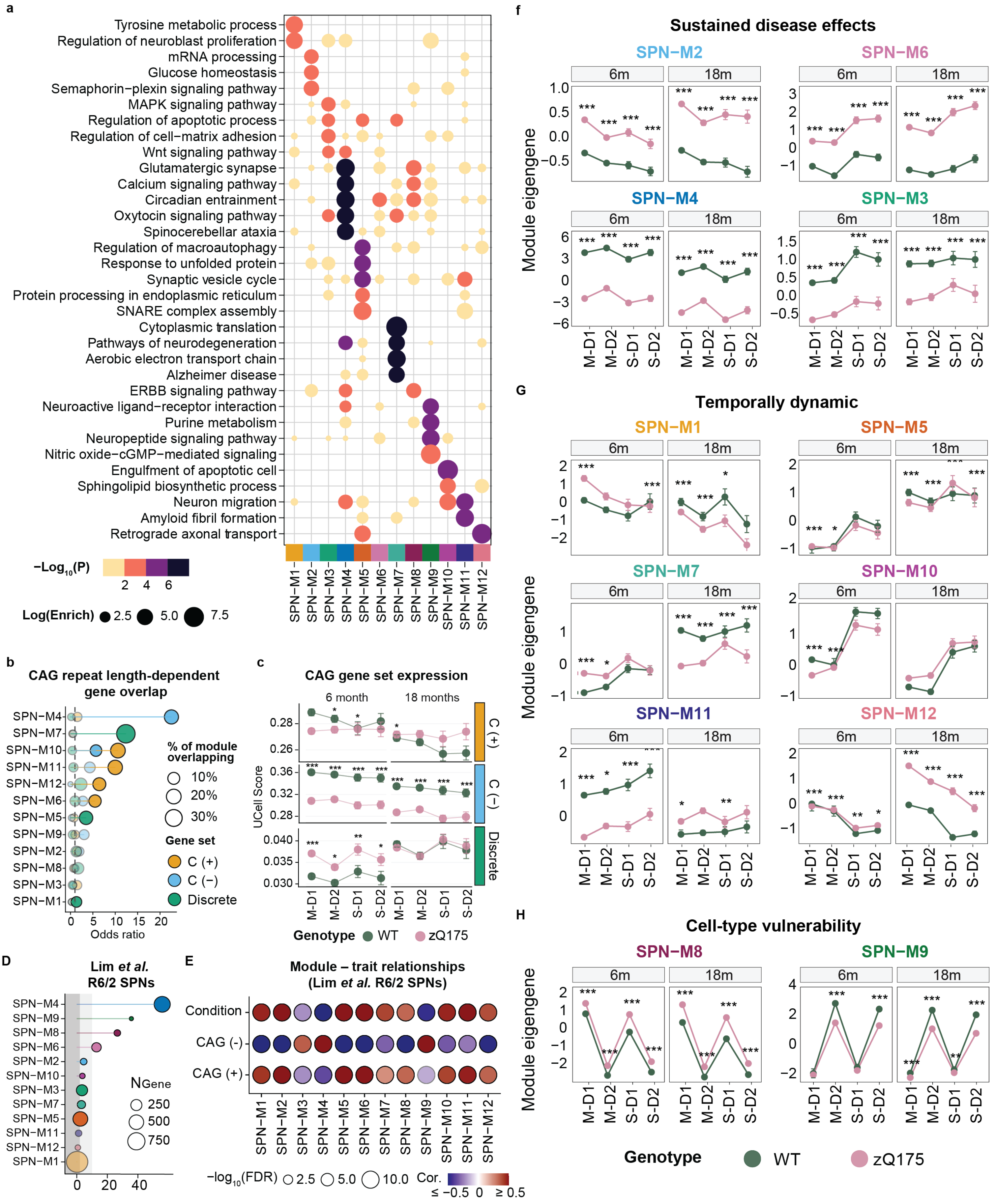

a
Tyrosine metabolic process
Regulation of neuroblast proliferation
mRNA processing
Glucose homeostasis
Semaphorin−plexin signaling pathway
MAPK signaling pathway
Regulation of apoptotic process
Regulation of cell−matrix adhesion
Wnt signaling pathway
Glutamatergic synapse
Calcium signaling pathway
Circadian entrainment
Oxytocin signaling pathway
Spinocerebellar ataxia
Regulation of macroautophagy
Response to unfolded protein
Synaptic vesicle cycle
Protein processing in endoplasmic reticulum
SNARE complex assembly
Cytoplasmic translation
Pathways of neurodegeneration
Aerobic electron transport chain
Alzheimer disease
ERBB signaling pathway
Neuroactive ligand−receptor interaction
Purine metabolism
Neuropeptide signaling pathway
Nitric oxide−cGMP−mediated signaling
Engulfment of apoptotic cell
Sphingolipid biosynthetic process
Neuron migration
Amyloid fibril formation
Retrograde axonal transport
−Log10(P)
2 4 6
Log(Enrich) 2.5 5.0 7.5
SPN−M1 SPN−M2 SPN−M3 SPN−M4 SPN−M5 SPN−M6 SPN−M7 SPN−M8 SPN−M9 SPN−M10 SPN−M11 SPN−M12
b
CAG repeat length-dependent gene overlap
SPN−M4 SPN−M7 SPN−M10 SPN−M11 SPN−M12 SPN−M6 SPN−M5 SPN−M9 SPN−M2 SPN−M8 SPN−M3 SPN−M1
0 5 10 15 20
Odds ratio
% of module overlapping
10%
20%
30%
Gene set
C (+)
C (−)
Discrete
c
CAG gene set expression
6 month
18 months
UCell Score
C (+)
C (−)
Discrete
M−D1 M−D2 S−D1 S−D2
Genotype WT zQ175
D
Lim et al. R6/2 SPNs
SPN−M4 SPN−M9 SPN−M8 SPN−M6 SPN−M2 SPN−M10 SPN−M3 SPN−M7 SPN−M5 SPN−M11 SPN−M12 SPN−M1
0 20 40
NGene
250
500
750
E
Module – trait relationships (Lim et al. R6/2 SPNs)
Condition
CAG (-)
CAG (+)
−log10(FDR) 2.5 5.0 10.0
Cor. ≤ −0.5 0 ≥ 0.5
f
Sustained disease effects
SPN−M2
SPN−M6
SPN−M4
SPN−M3
6m
18m
Module eigengene
M−D1 M−D2 S−D1 S−D2
G
Temporally dynamic
SPN−M1
SPN−M5
SPN−M7
SPN−M10
SPN−M11
SPN−M12
H
Cell-type vulnerability
SPN−M8
SPN−M9
Genotype WT zQ175

Supplementary Figure 4**. SPN gene co-expression gene networks capture temporal and subclass-specific expression patterns. a** Dot plot of select GO and KEGG pathway enrichment distinguishing co-expression modules. **b** Overlap of SPN hdWGCNA modules with CAG repeat-length dependent gene signatures in SPNs with > 150 CAG repeats, described in Handsaker et al. Odds ratio is shown on the x-axis. Dot color indicates gene set and size indicates the percent overlap of each gene set and the corresponding module genes. C (+): 262 genes positively correlation with CAG repeat length; CAG (-): 189 genes negatively correlation with CAG repeat length; D: 570 genes with discrete changes in SPNs. **c** CAG repeat length gene signature expression patterns in WT and zQ175 SPNs. Gene set scores were calculated by UCell and displayed separately for each SPN subtype by pathway (D1, dSPN; D2, iSPN) and compartment (M, Matrix; S, Striosome). Error bars indicate 95% confidence intervals for the average values. A linear model was fit for each subtype and timepoint, accounting for covariates of library batch and sex. *$p < 0.05$, **$p < 0.01$, ***$p < 0.001$. **d** Analysis of SPN hdWGCNA module preservation in a R6/2 mouse single-cell striatum dataset, Lim et al.[1] $Z > 10$, highly preserved; $Z > 2$, moderately preserved, and $Z < 2$, not preserved. **e** Module-trait relationships represented by the Pearson correlation for each module eigengene and covariate in the Lim et al. R6/2 SPN dataset. CAG (+) and CAG (-) scores were obtained by calculating the composite expression scores for each gene set with the UCell algorithm. % Mito: Mitochondrial RNA reads. **f-h** Module eigengene expression patterns in WT and zQ175 SPNs for each SPN subtype by pathway (D1, dSPN; D2, iSPN) and compartment (M, Matrix; S, Striosome). Error bars indicate 95% confidence intervals for the average values. A linear model was fit for each subtype and timepoint, accounting for covariates of library batch and sex. *$p < 0.05$, **$p < 0.001$, ***$p < 0.0001$.

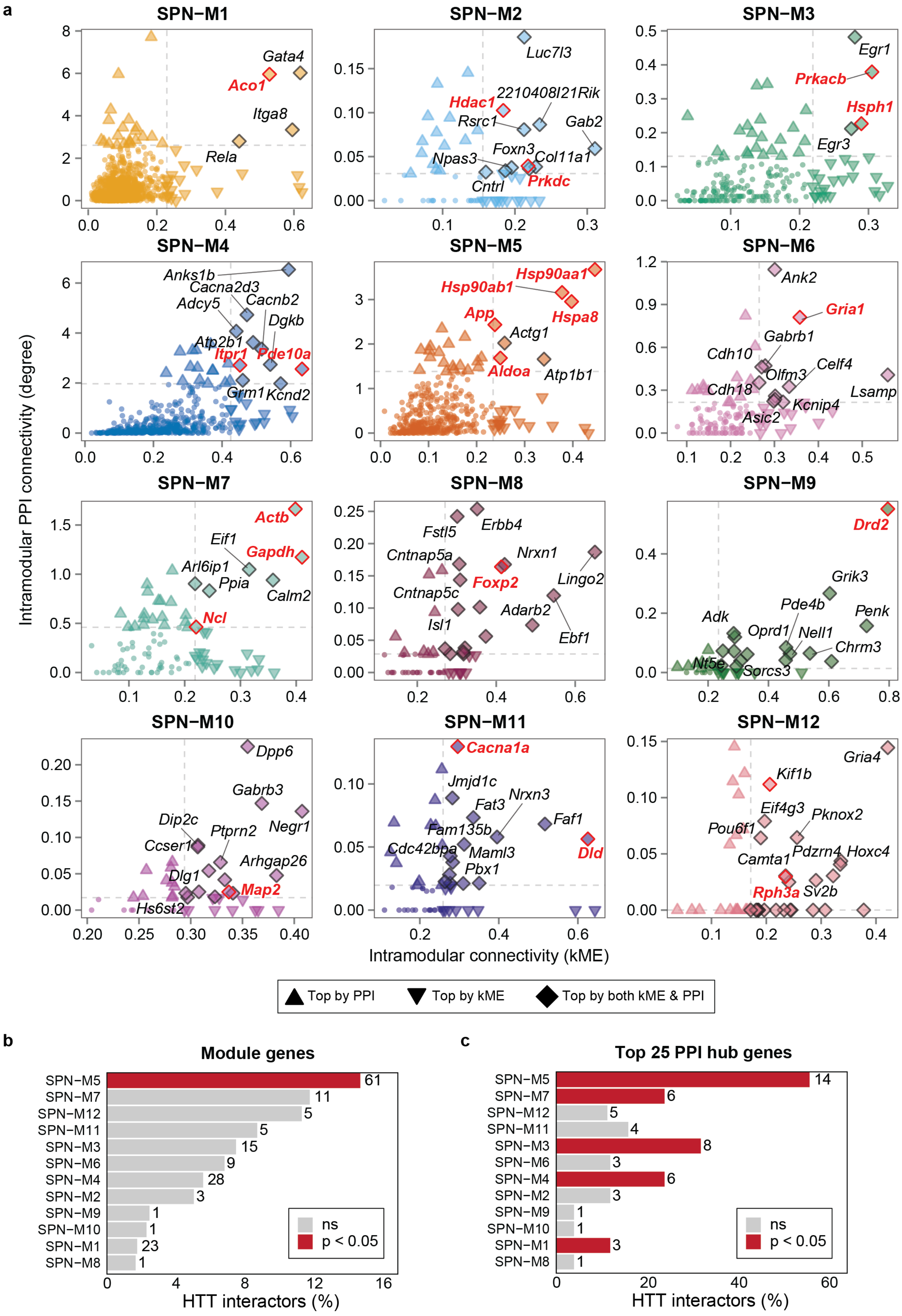
a
SPN−M1
Gata4
Aco1
Itga8
Rela
SPN−M2
Luc7l3
Hdac1
2210408I21Rik
Rsrc1
Gab2
Foxn3
Col11a1
Npas3
Cntrl
Prkdc
SPN−M3
Egr1
Prkacb
Hsph1
Egr3
SPN−M4
Anks1b
Cacna2d3
Cacnb2
Adcy5
Dgkb
Atp2b1
Itpr1
Pde10a
Grm1
Kcnd2
SPN−M5
Hsp90aa1
Hsp90ab1
Hspa8
App
Actg1
Aldoa
Atp1b1
SPN−M6
Ank2
Gria1
Gabrb1
Cdh10
Celf4
Olfm3
Cdh18
Lsamp
Kcnip4
Asic2
SPN−M7
Actb
Eif1
Gapdh
Arl6ip1
Ppia
Calm2
Ncl
SPN−M8
Erbb4
Fstl5
Cntnap5a
Nrxn1
Foxp2
Lingo2
Cntnap5c
Adarb2
Isl1
Ebf1
SPN−M9
Drd2
Grik3
Pde4b
Penk
Adk
Oprd1
Nell1
Chrm3
Nt5e
Sorcs3
SPN−M10
Dpp6
Gabrb3
Negr1
Dip2c
Ptprn2
Ccser1
Arhgap26
Dlg1
Map2
Hs6st2
SPN−M11
Cacna1a
Jmjd1c
Nrxn3
Fat3
Faf1
Fam135b
Cdc42bpa
Maml3
Dld
Pbx1
SPN−M12
Gria4
Kif1b
Eif4g3
Pknox2
Pou6f1
Camta1
Pdzrn4
Hoxc4
Rph3a
Sv2b
Intramodular PPI connectivity (degree)
Intramodular connectivity (kME)
Top by PPI
Top by kME
Top by both kME & PPI
b
Module genes
SPN−M5 61
SPN−M7 11
SPN−M12 5
SPN−M11 5
SPN−M3 15
SPN−M6 9
SPN−M4 28
SPN−M2 3
SPN−M9 1
SPN−M10 1
SPN−M1 23
SPN−M8 1
ns
p < 0.05
HTT interactors (%)
c
Top 25 PPI hub genes
SPN−M5 14
SPN−M7 6
SPN−M12 5
SPN−M11 4
SPN−M3 8
SPN−M6 3
SPN−M4 6
SPN−M2 3
SPN−M9 1
SPN−M10 1
SPN−M1 3
SPN−M8 1
ns
p < 0.05
HTT interactors (%)

Supplementary Figure 5**. Integration of functional protein association networks with SPN gene modules. a** Scatter plots of individual genes by module membership. The x-axis plots each module gene by eigengene-based connectivity (kME) which was used to determine the top hub genes in each module. The y-axis shows the degree of intramodular protein-protein interaction (PPI) connectivity for each gene within the gene network. Top hub genes in the integrated networks were defined as intersection of the top 25 genes by each ranking (kME and PPI degree). The shape of each dot represents the classification of the module genes as either top hits by kME, PPI or both. The top 10 shared top hub genes are annotated. **b** Enrichment of HTT protein interactors in all module genes and **c** in the top 25 hub genes by PPI degree, as defined by STRING database (Fisher's exact test, $p < 0.05$; Benjamini-Hochberg correction for multiple comparisons)

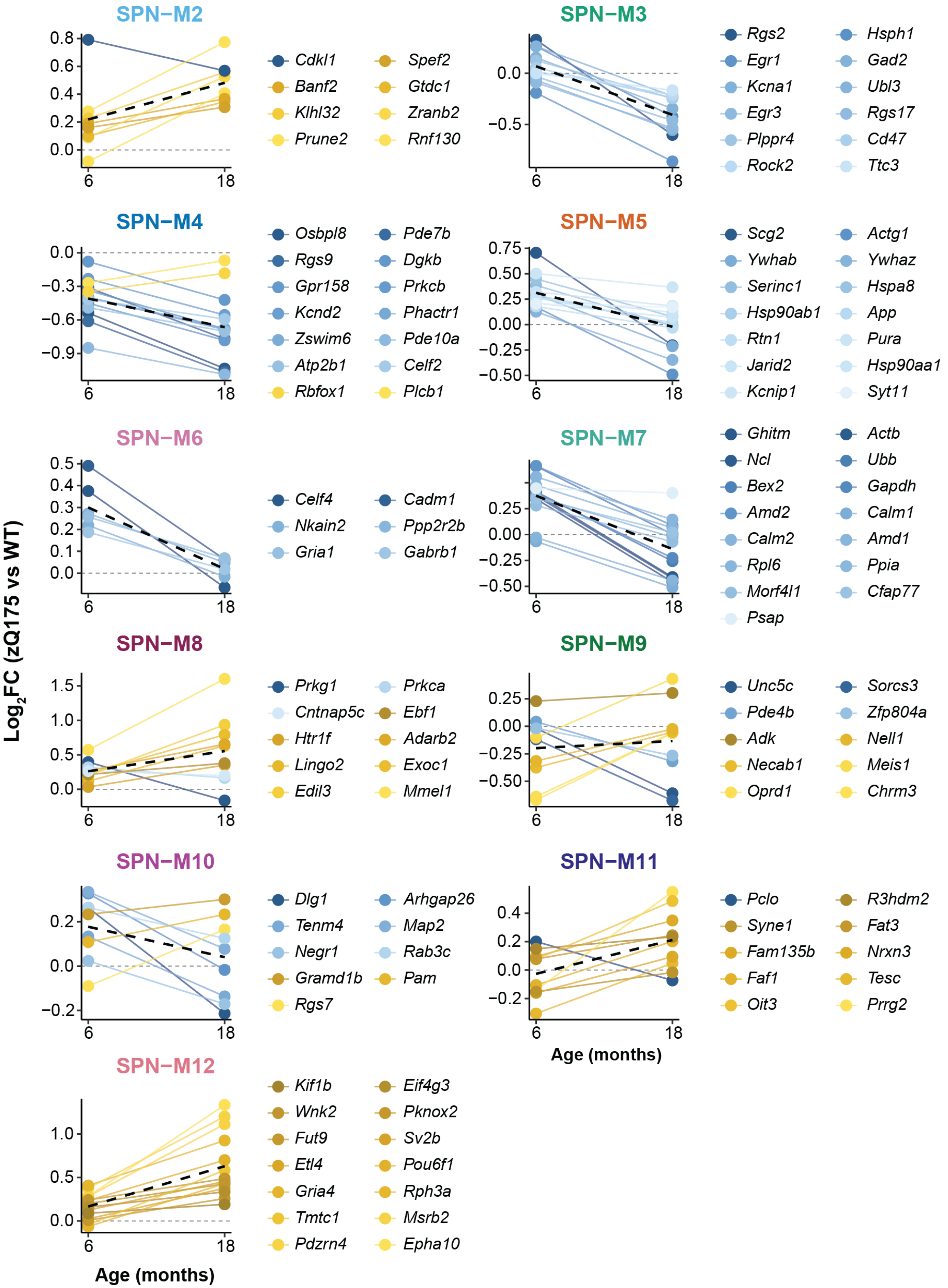
Log2FC (zQ175 vs WT)
Age (months)
SPN−M2
Cdkl1
Banf2
Klhl32
Prune2
Spef2
Gtdc1
Zranb2
Rnf130
SPN−M3
Rgs2
Egr1
Kcna1
Egr3
Plppr4
Rock2
Hsph1
Gad2
Ubl3
Rgs17
Cd47
Ttc3
SPN−M4
Osbpl8
Rgs9
Gpr158
Kcnd2
Zswim6
Atp2b1
Rbfox1
Pde7b
Dgkb
Prkcb
Phactr1
Pde10a
Celf2
Plcb1
SPN−M5
Scg2
Ywhab
Serinc1
Hsp90ab1
Rtn1
Jarid2
Kcnip1
Actg1
Ywhaz
Hspa8
App
Pura
Hsp90aa1
Syt11
SPN−M6
Celf4
Nkain2
Gria1
Cadm1
Ppp2r2b
Gabrb1
SPN−M7
Ghitm
Ncl
Bex2
Amd2
Calm2
Rpl6
Morf4l1
Psap
Actb
Ubb
Gapdh
Calm1
Amd1
Ppia
Cfap77
SPN−M8
Prkg1
Cntnap5c
Htr1f
Lingo2
Edil3
Prkca
Ebf1
Adarb2
Exoc1
Mmel1
SPN−M9
Unc5c
Pde4b
Adk
Necab1
Oprd1
Sorcs3
Zfp804a
Nell1
Meis1
Chrm3
SPN−M10
Dlg1
Tenm4
Negr1
Gramd1b
Rgs7
Arhgap26
Map2
Rab3c
Pam
SPN−M11
Pclo
Syne1
Fam135b
Faf1
Oit3
R3hdm2
Fat3
Nrxn3
Tesc
Prrg2
SPN−M12
Kif1b
Wnk2
Fut9
Etl4
Gria4
Tmtc1
Pdzrn4
Eif4g3
Pknox2
Sv2b
Pou6f1
Rph3a
Msrb2
Epha10

Supplementary Figure 6. **Temporal expression changes of SPN hdWGCNA hub genes reveal module-specific dynamics in HD.** Hub gene expression in SPNs between 6 and 18 months in zQ175 mice. Each panel displays the top hub genes by co-expression module. The top 25 hub genes were filtered by expression pattern (progressive, early and late), genes with no change from WT between timepoints are excluded from visualization. Lines connect genes between time points and color is by interaction effect. FDR < 0.05 and $|\log_2$ fold change, zQ175 vs. WT$| \geq 0.2$ or $|\log_2$ fold change, interaction$| \geq 0.2$.

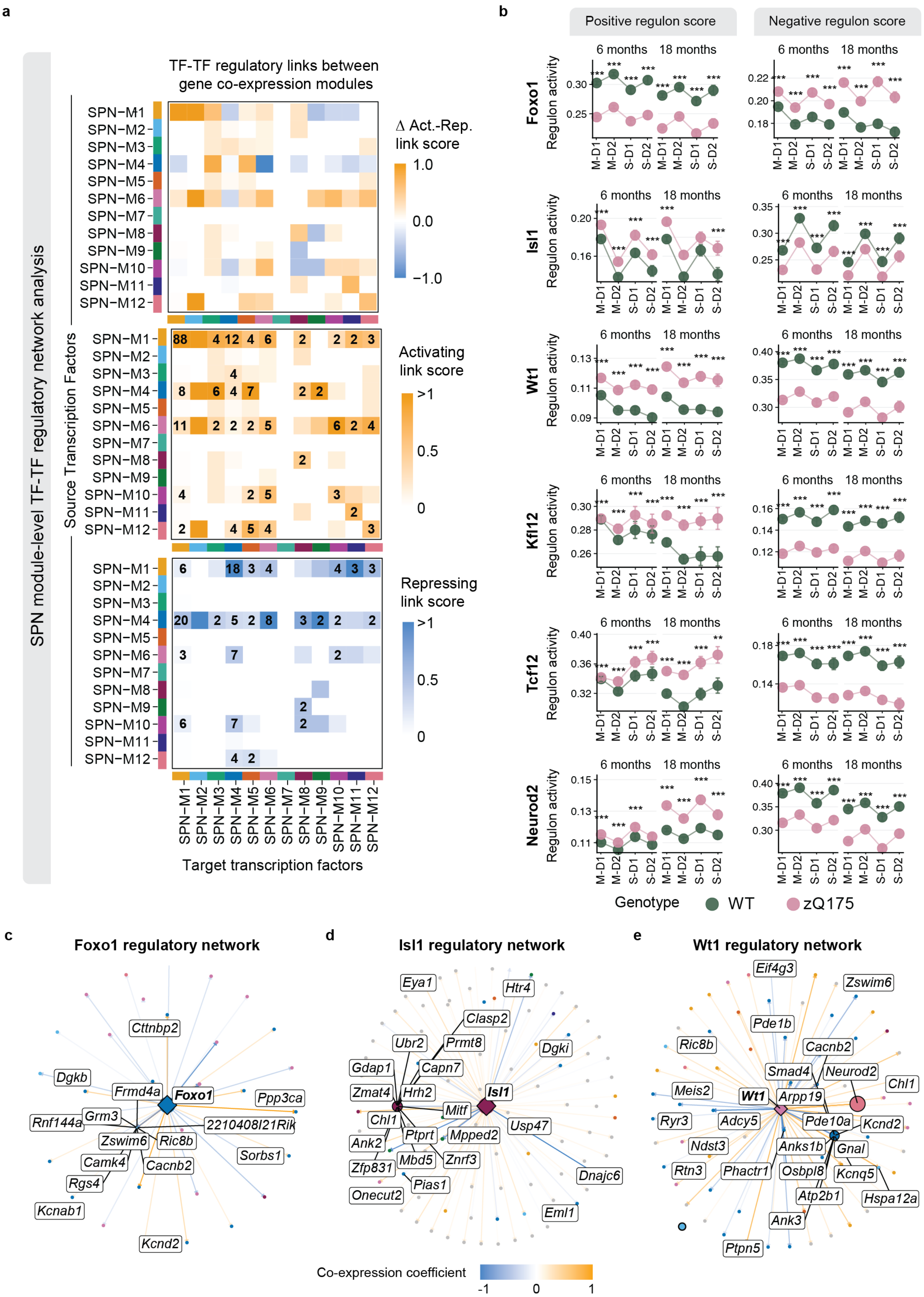
a
SPN module-level TF-TF regulatory network analysis
TF-TF regulatory links between gene co-expression modules
Δ Act.-Rep. link score
1.0
0.0
−1.0
Activating link score
>1
0.5
0
Repressing link score
>1
0.5
0
Source Transcription Factors
SPN−M1
SPN−M2
SPN−M3
SPN−M4
SPN−M5
SPN−M6
SPN−M7
SPN−M8
SPN−M9
SPN−M10
SPN−M11
SPN−M12
Target transcription factors
b
Positive regulon score
Negative regulon score
6 months
18 months
Regulon activity
Foxo1
Isl1
Wt1
Klf12
Tcf12
Neurod2
M-D1
M-D2
S-D1
S-D2
Genotype
WT
zQ175
c
Foxo1 regulatory network
Foxo1
Cttnbp2
Dgkb
Frmd4a
Ppp3ca
Rnf144a
Grm3
2210408I21Rik
Zswim6
Ric8b
Sorbs1
Camk4
Cacnb2
Rgs4
Kcnab1
Kcnd2
d
Isl1 regulatory network
Isl1
Eya1
Htr4
Clasp2
Ubr2
Prmt8
Dgki
Gdap1
Capn7
Zmat4
Hrh2
Mitf
Chl1
Usp47
Ptprt
Mpped2
Ank2
Mbd5
Znrf3
Zfp831
Dnajc6
Pias1
Onecut2
Eml1
e
Wt1 regulatory network
Wt1
Eif4g3
Zswim6
Pde1b
Ric8b
Cacnb2
Smad4
Neurod2
Chl1
Meis2
Arpp19
Ryr3
Adcy5
Pde10a
Kcnd2
Ndst3
Anks1b
Gnal
Rtn3
Phactr1
Osbpl8
Kcnq5
Atp2b1
Hspa12a
Ank3
Ptpn5
Co-expression coefficient
-1
0
1

Supplementary Figure 7**. Analysis of module and TF regulatory networks in striatal projection neurons.**

**a** Heatmap representation of TF-TF regulatory connections. Regulatory scores represent the number of TF-gene links the source to target module, normalized by the total number of TFs in the source module. Top panel shows the difference between the positive and negative regulatory score. A positive score suggests a module is activating, whereas a negative score indicates repressive activity. Middle panel shows only activating links and the bottom panel shows repressive links. **b** Mean regulon activity scores across SPN subtypes at 6 and 18-months, compared between zQ175 and WT. Error bars indicate 95% confidence intervals for the average values. A linear model was fit for each subtype and timepoint, accounting for covariates of library batch and sex. *$p < 0.05$, **$p < 0.001$, ***$p < 0.0001$. **c-e** Regulatory network plots for select TFs showing all predicted target genes as nodes. Labeled nodes represent CAG repeat-length dependent genes as defined in Handsaker et al. Nodes are shown only for genes with a Pearson co-expression correlation coefficient > 0.15. Only primary targets are displayed for each TF in the network. Direct TF targets are represented by highlighted nodes shaped as circles.

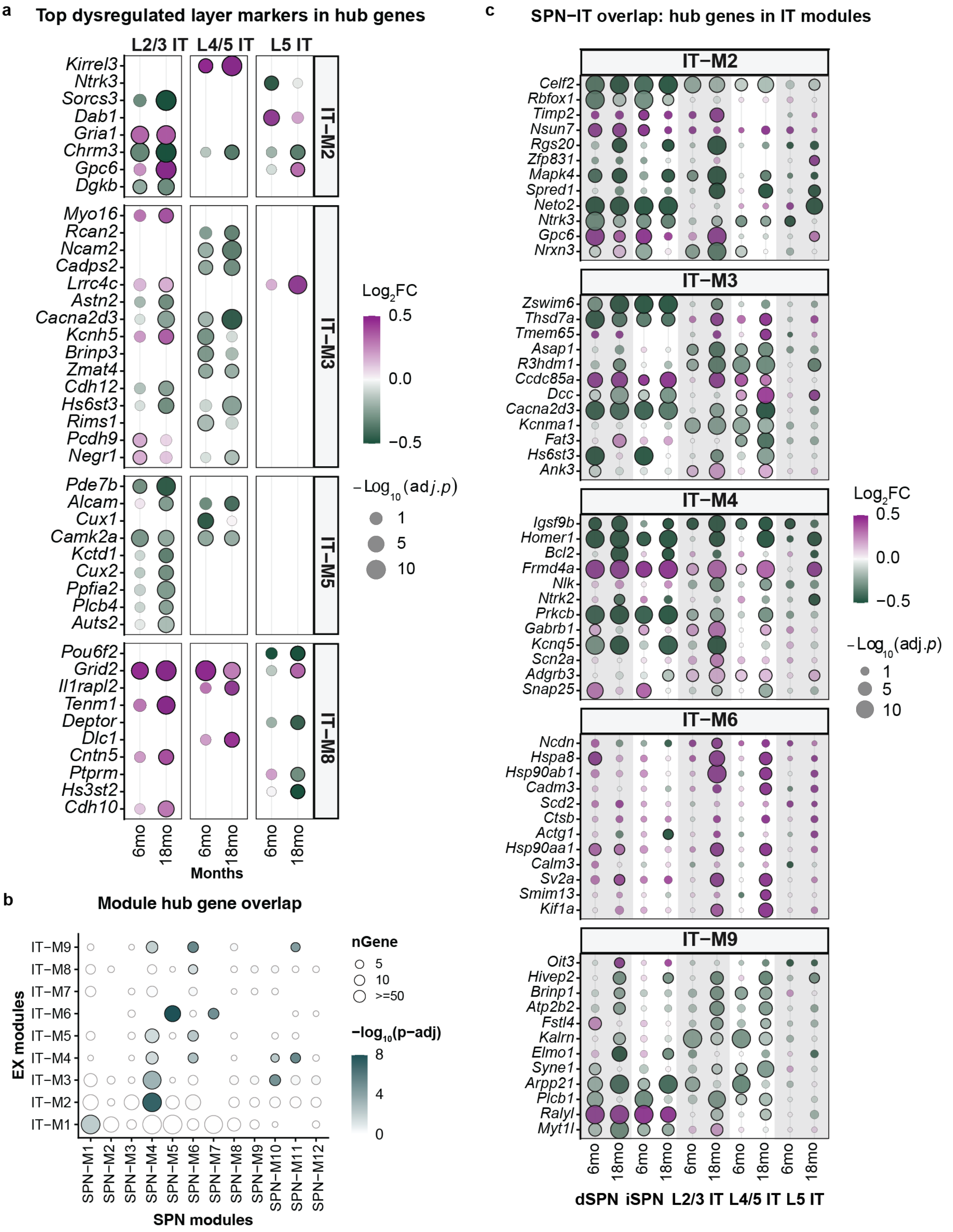

a
Top dysregulated layer markers in hub genes
L2/3 IT
L4/5 IT
L5 IT
IT−M2
Kirrel3
Ntrk3
Sorcs3
Dab1
Gria1
Chrm3
Gpc6
Dgkb
IT−M3
Myo16
Rcan2
Ncam2
Cadps2
Lrrc4c
Astn2
Cacna2d3
Kcnh5
Brinp3
Zmat4
Cdh12
Hs6st3
Rims1
Pcdh9
Negr1
IT−M5
Pde7b
Alcam
Cux1
Camk2a
Kctd1
Cux2
Ppfia2
Plcb4
Auts2
IT−M8
Pou6f2
Grid2
Il1rapl2
Tenm1
Deptor
Dlc1
Cntn5
Ptprm
Hs3st2
Cdh10
6mo
18mo
Months
Log2FC
0.5
0.0
−0.5
−Log10(adj.p)
1
5
10
b
Module hub gene overlap
EX modules
IT−M9
IT−M8
IT−M7
IT−M6
IT−M5
IT−M4
IT−M3
IT−M2
IT−M1
SPN−M1
SPN−M2
SPN−M3
SPN−M4
SPN−M5
SPN−M6
SPN−M7
SPN−M8
SPN−M9
SPN−M10
SPN−M11
SPN−M12
SPN modules
nGene
5
10
>=50
−log10(p−adj)
8
4
0
c
SPN−IT overlap: hub genes in IT modules
IT−M2
Celf2
Rbfox1
Timp2
Nsun7
Rgs20
Zfp831
Mapk4
Spred1
Neto2
Ntrk3
Gpc6
Nrxn3
IT−M3
Zswim6
Thsd7a
Tmem65
Asap1
R3hdm1
Ccdc85a
Dcc
Cacna2d3
Kcnma1
Fat3
Hs6st3
Ank3
IT−M4
Igsf9b
Homer1
Bcl2
Frmd4a
Nlk
Ntrk2
Prkcb
Gabrb1
Kcnq5
Scn2a
Adgrb3
Snap25
IT−M6
Ncdn
Hspa8
Hsp90ab1
Cadm3
Scd2
Ctsb
Actg1
Hsp90aa1
Calm3
Sv2a
Smim13
Kif1a
IT−M9
Oit3
Hivep2
Brinp1
Atp2b2
Fstl4
Kalrn
Elmo1
Syne1
Arpp21
Plcb1
Ralyl
Myt1l
6mo
18mo
dSPN
iSPN
L2/3 IT
L4/5 IT
L5 IT
Log2FC
0.5
0.0
−0.5
−Log10(adj.p)
1
5
10

Supplementary Figure 8. **IT neuron layer-specific and core disease signatures represented in module hub genes. a** Differentially expressed layer-specific marker genes in top hub genes for cortical layer enriched modules (IT-M2, M3, M5 and M6), separately for IT subclusters defined by unsupervised clustering. **b** Module gene overlap analysis between IT and SPN hdWGCNA modules. **c** Differential expression of select top shared hub genes between IT and SPN modules, grouped by module assignment in IT neurons.
